# Rapid centromere turnover and the adaptive radiation of lemurs

**DOI:** 10.64898/2026.05.16.725662

**Authors:** Mihir Trivedi, Francesca Gianfrate, Luciana de Gennaro, Marcelo Ayllon, Katherine M. Munson, Kendra Hoekzema, DongAhn Yoo, Erin E. Ehmke, Anne D. Yoder, Stephen Chang, Chinmay Lalgudi, Mark Krasnow, Mario Ventura, Evan E. Eichler

## Abstract

Centromeres represent essential chromosomal structures required for faithful chromosome segregation during cell division but are paradoxically hypermutable, leading to centromere drive and reproductive isolation in closely related species. Using long-read sequencing, we generate nearly complete genomes (2.1-2.5 Gbp) from eight lemur species and characterize the sequence, epigenetic and cytogenetic structure of 223 strepsirrhini centromeres providing an alternative primate perspective of centromere evolution. No lemur centromere consists of α-satellite DNA that typifies the haplorhine lineage; instead, each species evolved its own distinct higher-order centromeric repeat sequence, varying substantially in both monomer length (ranging from 41-548 bp) and primary sequence composition (GC percentages 28.7-67.9%) including centromere cooption of telomeric repeats in brown lemurs. Most centromeres show characteristic hypomethylation dip regions (110-300 kbp) as candidates for kinetochore attachment. The centromere sequence motif shows no apparent sequence homology among lemur genera, even for species separated by less than 15 million years (*Lemur* and *Eulemur*). We estimate a >6-fold increased rate in primary centromeric motif turnover in strepsirrhines when compared to haplorhines and this occurred in conjunction with positive selection of the CENP-B protein in lemur lineages. We propose that lemur radiation and centromere diversification are linked, whereby accelerated motif turnover provides a stasipatric barrier contributing to rapid chromosomal evolution.

## INTRODUCTION

At the most basal level, the primate order is classified into two clades: strepsirrhines and haplorhines. The suborder Strepsirrhini is further comprised of lemurs (Lemuriformes), lorises (Lorisidae), and galagos (Galagidae) (Fleagle, 2013). Lemuriformes have been a focus of primate research due to their diversity and a restricted geographical distribution, which is limited to the island of Madagascar (Yoder, 2007). With some estimates of species diversity ranging as high as 112 extant and 17 extinct species concentrated in less than 15% of the planet’s land area, lemurs are uniquely well suited to provide insights into animal speciation (Mittermeier et al., 2022; Godfrey et al., 2010). They are thought to have diverged from the clade containing Old World and New World haplorhine primates 70–79 million years ago and represent the deepest split among the primates, rendering them critical to understand early primate adaptations (Horvath & Willard, 2007; dos Reis et al., 2018).

Owing to this complicated diversification, the evolutionary history of lemurs and the forces that shaped their speciation remain a matter of debate. A wealth of karyotypic data has been gathered over the last 50 years for all major lineages within the Lemuriformes, including the Lemuridae, Indriidae, Cheirogaleidae, and monotypic aye-aye, *Daubentonia* (Rumpler & Dutrillaux, 1976; Rumpler & Dutrillaux, 1979; Rumpler et al., 1983; Rumpler et al., 1988). Ultimately, this work culminated with informed speculation on the ancestral karyotype for the lemuriform clade as well as the sequence of chromosomal fissions and fusions that produced the lineage-specific karyotypes of living lemurs (Warter et al., 2005). While lemur phylogeny and evolution are still an active area of research (Everson et al., 2025a; Orkin et al., 2025), it is largely agreed that Madagascar’s environmental heterogeneity must have provided a variety of niches for ancestral lemur species to speciate after their oceanic dispersal from mainland Africa (Samonds et al., 2012; Yoder et al., 1996).

Despite their central role in understanding primate evolution, there is a paucity of high-quality genomes, and regions such as centromeres have largely been uncharacterized. Among the publicly available reference quality lemur genomes from 41 species, 38 are derived from Illumina short reads (Guevara et al., 2021; Kuderna et al., 2023), while three recent genomes incorporate PacBio and/or Oxford Nanopore Technologies (ONT) long reads (Everson et al., 2025b; Palmada-Flores et al., 2022; Versoza & Pfeifer, 2024). Most of these are highly fragmented, with NCBI labeling seven assemblies as “chromosome level.” Some lemur genomes remain unpublished with preliminary analysis done either independently or as a part of sequencing consortia (Supplementary Table 1). Many of these early assemblies are limited by their low contiguity and high numbers of unplaced scaffolds, rendering them insufficient for the study of centromere structure and evolution.

Recent advances in deep long-read sequencing from both ultra-long ONT as well as PacBio high-fidelity (HiFi) coupled to algorithmic improvements (Cheng et al., 2021), such as the Verkko assembler (Antipov et al., 2025), have resulted in much longer contiguous assemblies traversing some of the longest repeat repetitive regions of primate genomes (Nurk et al., 2022), including centromeres (Logsdon et al., 2024; Logsdon et al., 2025). In order to systematically characterize centromeres among the Strepsirrhini, we focused on the generation of *ab initio* long-read assemblies of eight species. We use these new assemblies to focus on the genetic characterization of lemur centromeres. To date, only the gray mouse lemur and the aye-aye centromeres have been characterized (Larsen et al., 2017; Lee et al., 2011). The former centromere consists of a 53 bp motif (Mm53) that is tandemly repeated to create complex arrays from 400 kbp to 3.2 Mbp, depending on the chromosome. The aye-aye was reported to have two consensus sequences of 146 bp and 268 bp. We extend this work to a newer assembly of the gray mouse lemur and seven other lemuriforms representing 40 million years of diversity. Here, we describe differences in the primary repeat motifs and relative chromosomal location in the context of species-specific higher-order repeat (HOR) structure as well as epigenetic properties associated with the hypomethylation centromere dip region (CDR) thought to define the location of the kinetochore binding among humans and other primates (Logsdon et al., 2024; Yoo et al., 2025).

## RESULTS

### Lemur genome assembly and synteny breakpoints

We selected eight lemur species representing both short and long genetic distances over the last 40 million years of strepsirrhine evolution (Everson et al., 2025a): *Lemur catta* (LCA, ring-tailed lemur), *Microcebus murinus* (MMU, gray mouse lemur), *Cheirogaleus medius* (CME, fat-tailed dwarf lemur), *Propithecus coquereli* (PCO, Coquerel’s sifaka), *Eulemur collaris* (ECO, collared brown lemur), *Varecia rubra* (VRU, red ruffed lemur), *Varecia variegata* (VVA, black and white ruffed lemur), and *Daubentonia madagascariensis* (DMA, aye-aye) (Figure 1a). With the exception of *M. murinus*, which was assembled independently (Methods), we obtained primary blood lymphocytes for the other seven species from the Duke Lemur Center (DLC) and extracted high molecular weight DNA. The same material was sequenced with both long-read technologies, PacBio HiFi and ONT, to obtain high coverage for each species (median coverages of 64x and 34x, respectively [Table 1]). For the purpose of centromere reconstruction (often associated with Mbp of tandem repeats), we focused on the production of ultra-long ONT reads to effectively span such regions of the genomes.

**Figure 1.**
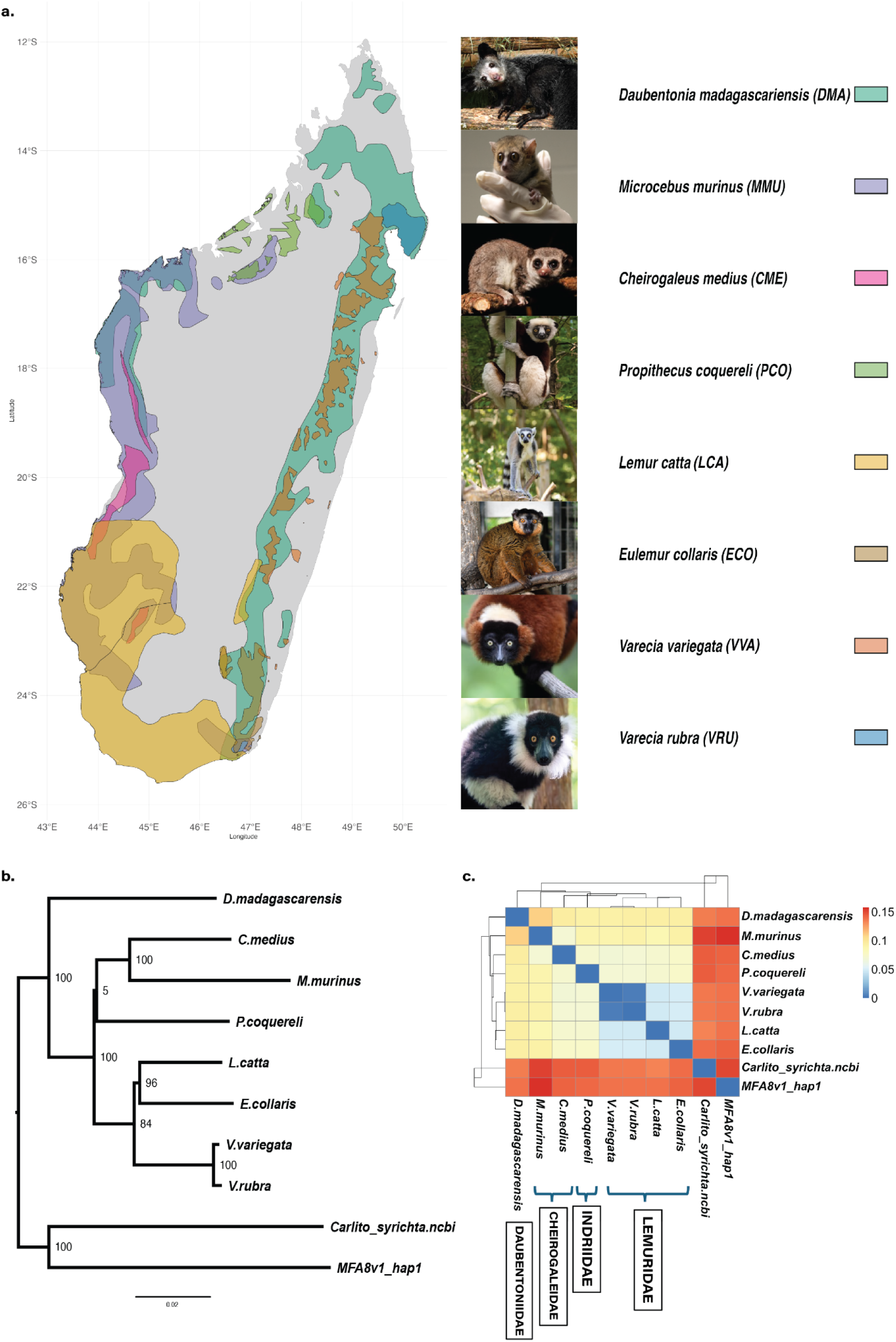
Lemur species and their phylogenetic relationships. a, The geographic distribution of lemur species, used in this study, on the island of Madagascar per the latest release of the International Union of Conservation of Nature (IUCN). The color scheme is described with scientific names and their three letter codes used in the paper. b, Phylogenetic relationship among species calculated based on genetic distances (Methods), using *Macaca fascicularis* and *Carlito syrichta* sequences as the outgroups. Bootstrap values are shown for each node. c, Heatmap showing the pairwise distances between species and delineating four primate families represented by the eight species.

**Table 1.**
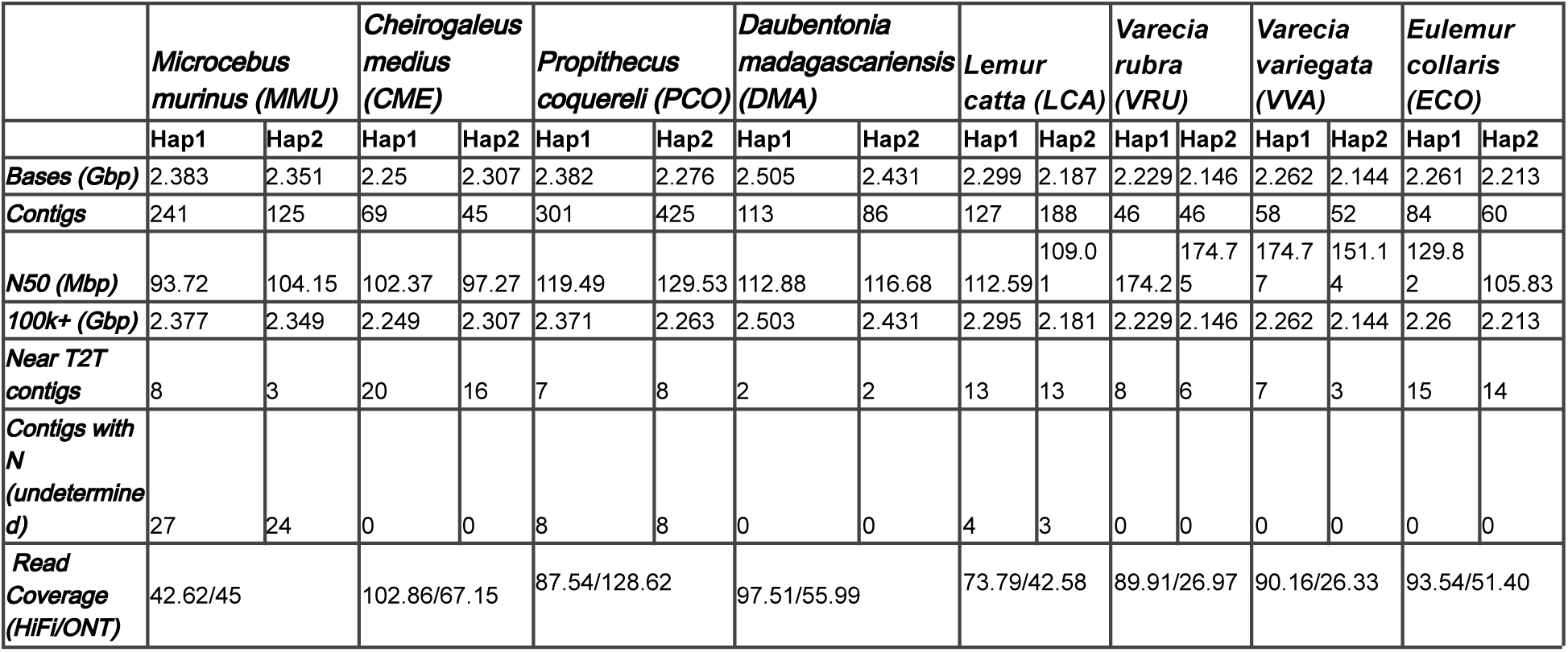
Summary of sequence and assembly statistics.

In general, the assemblies were generated without chromosomal-level phasing since our focus was to locally assemble centromeres, with two exceptions. For two species, *Lemur catta* (LCA) and *Propithecus coquereli* (PCO), we collected parental samples and phased the assembly with parental Illumina short-read whole-genome sequences. All non-parental phased assemblies were generated by hifiasm (version 0.19.9 or 0.23.0) (Cheng et al., 2021), while parentally phased assemblies were generated with both hifiasm and Verkko (2.1.2) (Antipov et al., 2025), and the best out of the two was chosen based on contiguity and other parameters. By using these techniques, we successfully generated highly contiguous assemblies, each featuring multiple near-telomere-to-telomere (T2T) contigs across all species studied (Supplementary Figure S1).

To confirm the genetic relationship among species, we computed the genetic distances among the genomes to construct a bootstrapped neighbor-joining tree using Mashtree (Katz et al., 2019; Ondov et al., 2016). For phylogenetic reconstruction, we selected the crab-eating macaque, *Macaca fascicularis* (MFA) T2T genome (Zhang et al., 2025), and the tarsier, *Carlito syrichta* (CSY) (Schmitz et al., 2016) genome, as outgroups. The phylogenetic tree supports the generally accepted phylogeny of the taxa, including the assignment of congeneric species (Everson et al., 2025a); however, the split between *Propithecus* and Cheirogaleidae (*Microcebus* and *Cheirogaleus*) was not confidently resolved as evidenced by the lower bootstrap support (Figure 1b). The unresolved nature of the *Propithecus* genus and the overall Indriidae family among other lemur families has been previously described (Harrera and Davalos, 2016). Aye-aye (*Daubentonia*) is the clear outgroup of all the lemur species, while true lemurs (Lemuridae) form the innermost cluster.

We also aligned the eight assembled strepsirrhine genomes to the finished human reference genome, T2T-CHM13, using Anchorwave aligner (Song et al., 2022) (Methods). Given their deep divergence, strepsirrhines are expected to have a large number of chromosomal rearrangements with respect to other haplorhine primates. With respect to humans, *Lemur catta* (LCA) has 159 synteny breakpoints. Five chromosomes from T2T-CHM13: 1, 7, 12, 22 and X, are fully conserved with respect to synteny, albeit with intrachromosomal rearrangements and changes in gene order (Figure 2a). As a control, we also performed a similar synteny analysis comparing LCA against the recently completed genome of MFA (Zhang et al., 2025) (Supplementary Figure S2).

**Figure 2.**
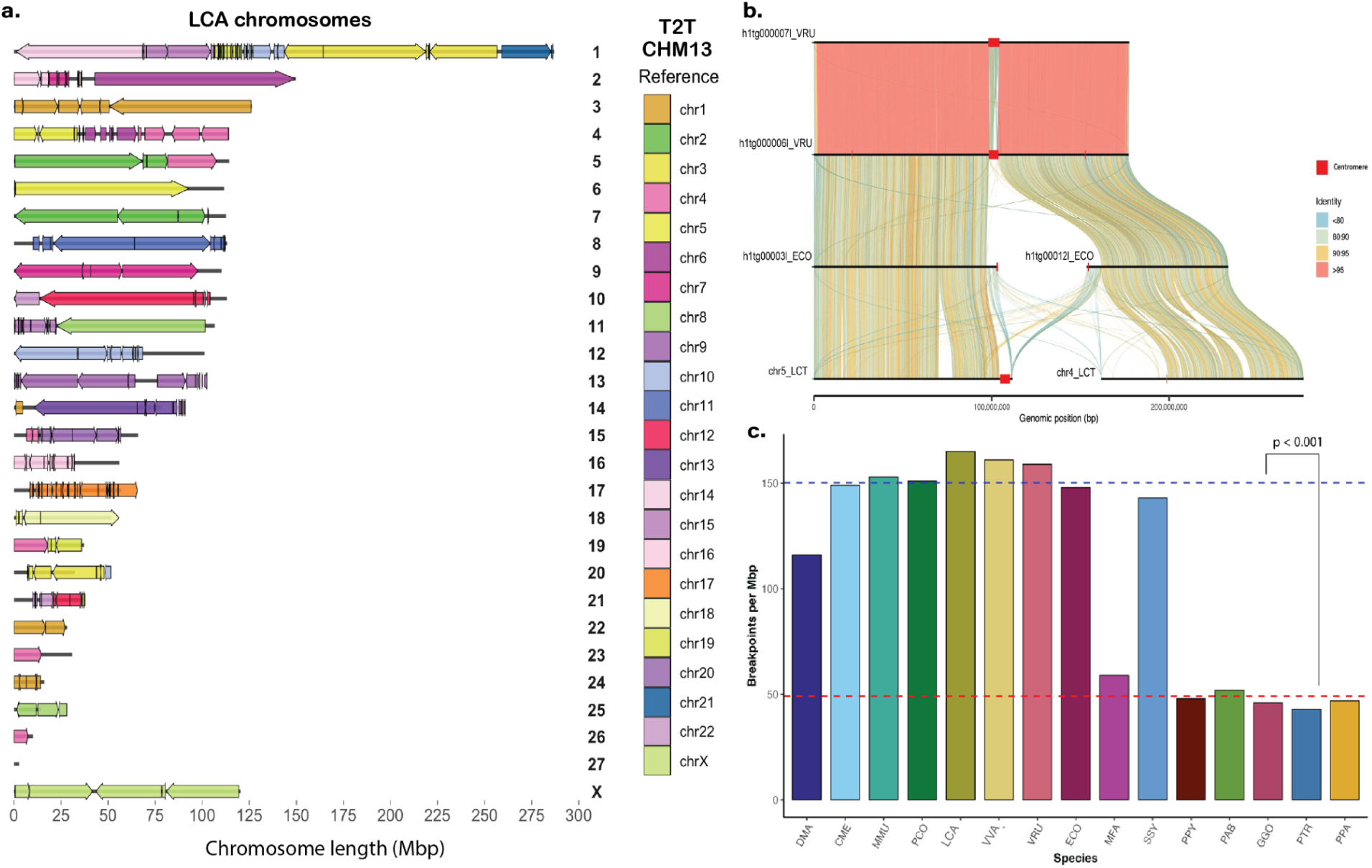
Chromosome level assembly and synteny breakpoints. a, Synteny between T2T-CHM13 and *Lemur catta* (LCA) chromosomes, with LCA on the left and human (T2T-CHM13) chromosome colors on the right. Arrows show the orientation of T2T-CHM13 synteny blocks and straight lines show the regions with no syntenic matches. b, Individual contig/chromosome level synteny with SVbyEye. Single contig from two sister species, VRU and VVA, aligns to two different contigs in ECO and two different chromosomes in LCA with breakpoints occurring over a centromere (red). The color scheme shows % identity between aligned sequences with >95% between VRU and VVA. c, Comparison of the total number of syntenic breakpoints of the lemur species in this study and nonhuman apes (Yoo et al., 2025) compared to human (T2T-CHM13). The mean number of non-human apes’ breakpoints (red) and lemur syntenic breakpoints (blue) is compared p < 0.001.

We consolidated breakpoint counts across three primate groups: eight strepsirrhines, eight Old World monkeys and apes, and the human T2T-CHM13 reference. A representative syntenic segment across four closely related species, LCA, ECO, VVA, and VRU, illustrates the disruption of sequence identity and collinearity among lineages that diverged approximately 15 million years ago (Figure 2b). As expected, we observe a significant difference (p<0.0001, Wilcoxon-Rank test) between number of breakpoints when comparing strepsirrhines and haplorhines to human or macaque (Figure 2c, Supplementary Figure S3). Overall, there is a gradual decline in the number of syntenic blocks as a function of genetic distance with the conspicuous exception of siamang (SSY), which is known to have undergone an exceptionally rapid karyotype evolution among the apes (Carbone et al., 2014, Yoo et al., 2025). The aye-aye (DMA) shows the lowest number of breakpoints when compared to complete haplorhine genomes consistent with its rather conserved karyotype of 2n = 30 (Poorman-Allen and Izzard, 1990).

### Lemur centromere motif discovery, validation and characterization

Since α-satellite motifs that are organized into HOR satellite arrays typically define centromeres in haplorhine primates (Yoo et al., 2025), we specifically searched within contiguous genome assemblies for the presence of long tandem repeats using Tandem Repeats Finder (TRF) (Benson, 1999). For each genome, we identified several potential candidates. Next, we designed a set of in-house scripts to then characterize a consensus repeat and used these to define potential higher-order structures on each chromosome as well as their distribution among chromosomes within each species. This is similar to methods used previously to define centromeres in mouse lemur and other mammal species (Melters et al., 2013; Larsen et al., 2017).

Employing this approach, we identified 223 potential centromeres from a total of ∼432 lemur contigs. Of note, not all chromosomes were sequenced completely and the karyotype for ECO is not precisely known. Nevertheless, we identified 10 distinct monomer motifs for seven strepsirrhine species (excluding Mm53), including both high GC and high AT-repeat motifs (Figure 6 and Table 2). We designated each using the convention previously applied to *Microcebus murinus*; namely, species acronym followed by motif length).

**Figure 3.**
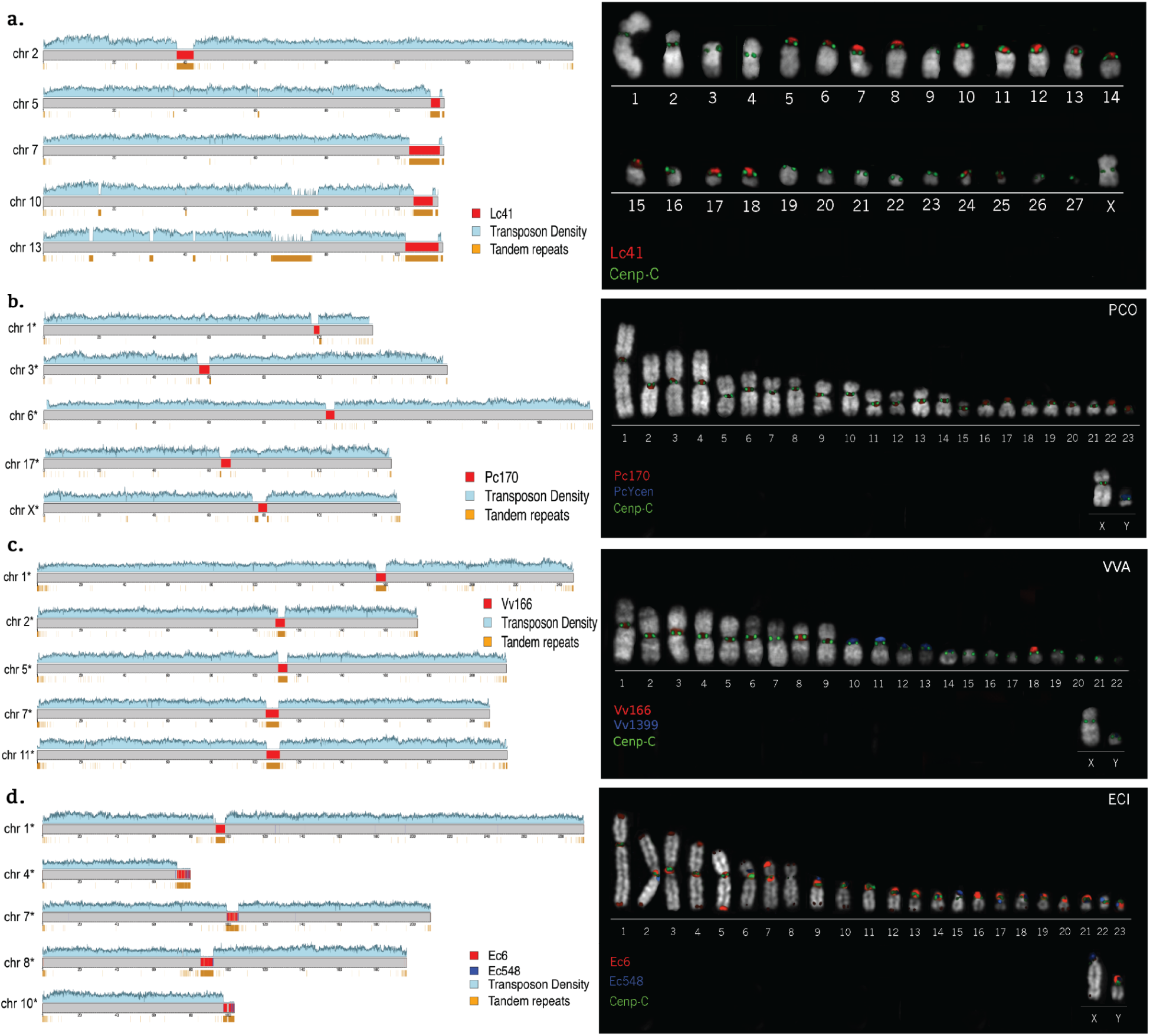
Centromere sequence validation by FISH. We depict five chromosome contig sequences (left) as ideograms of four lemur species: a. LCA, b. PCO, c. VVA, and d. ECO based on sequence and assembly. Centromeric (red) and pericentromeric repeats (blue for VVA and ECO) are shown with respect to transposon density, which drastically reduces at the putative centromeres. Co-immunohybridization (right panels) experiments showing FISH with candidate sequence probes (red) and CENP-C antibody signal (green) validate centromeric localization (closely related ECI is used in lieu of ECO). All chromosomes with an asterisk (*) are named according to their synteny with respect to LCA.

Overall, we find that centromeres exhibit distinct characteristics among diverse lemur species, with each genus possessing distinct single monomers or even two monomers in the same species. In general, the total average length of HORs appear smaller (typically less than 5 Mbp) in size when compared to eight haplorhine species that were recently T2T sequenced (Yoo et al., 2025; Zhang et al., 2025) (Figure 4). This difference is also statistically significant when we perform centromere group comparisons between strepsirrhine and haplorhine (Figure 4a, Supplementary Figure S4) (p < 2.22e-16, Wilcoxon rank sum test). We also compared monomer sequence divergence with respect to each consensus sequence (Table 2) and contrasted it with each other and canonical alpha-satellite divergence for haplorhines. Most lemur species follow a unimodal distribution: LCA, VRU, PCO, MMU, and CME (Figure 4b). DMA shows a distinct tri-modality in distribution as the centromeric repeats have diverged to give two new shorter “variants” of the 267 bp monomer, with consensus sizes of 222 and 209 bp. VVA shows a bimodal distribution, which points to other interspersed sequences in its centromeres.

Because we typically resolve two homologues for each chromosome, we could also compare the allelic and non-allelic sequence identity of centromeres and contrast it with unique flanking sequences in the assemblies. As expected, across species allelic centromeres share higher sequence identity to each other than to those from other chromosomes with identities ranging from lowest allelic median for VVA at 91.18 ± 3.94% to highest for MMU at 96.94 ± 3.17%. Non-allelic centromere identity medians ranged from 86.43 ± 1.22% again in VVA to 94.59 ± 1.76% in ECO. For each strepsirrhine species, we summarized the HOR structure using StainedGlass (Vollger et al., 2022) and HiCAT (Gao et al., 2023) and use the tool’s standard notation to describe HOR organization (i.e., R1L6 defines the highest-ranked HOR, which maximizes both coverage and repeat fidelity (Methods) while L denotes the length defined by the number of repeating monomer units in an HOR cassette). We also investigate 5-methyl cytosine features of the corresponding heterochromatin associated with each centromere, searching for evidence of a CDR (Figure 5). We summarize these centromere features individually for each species below.

**Figure 4.**
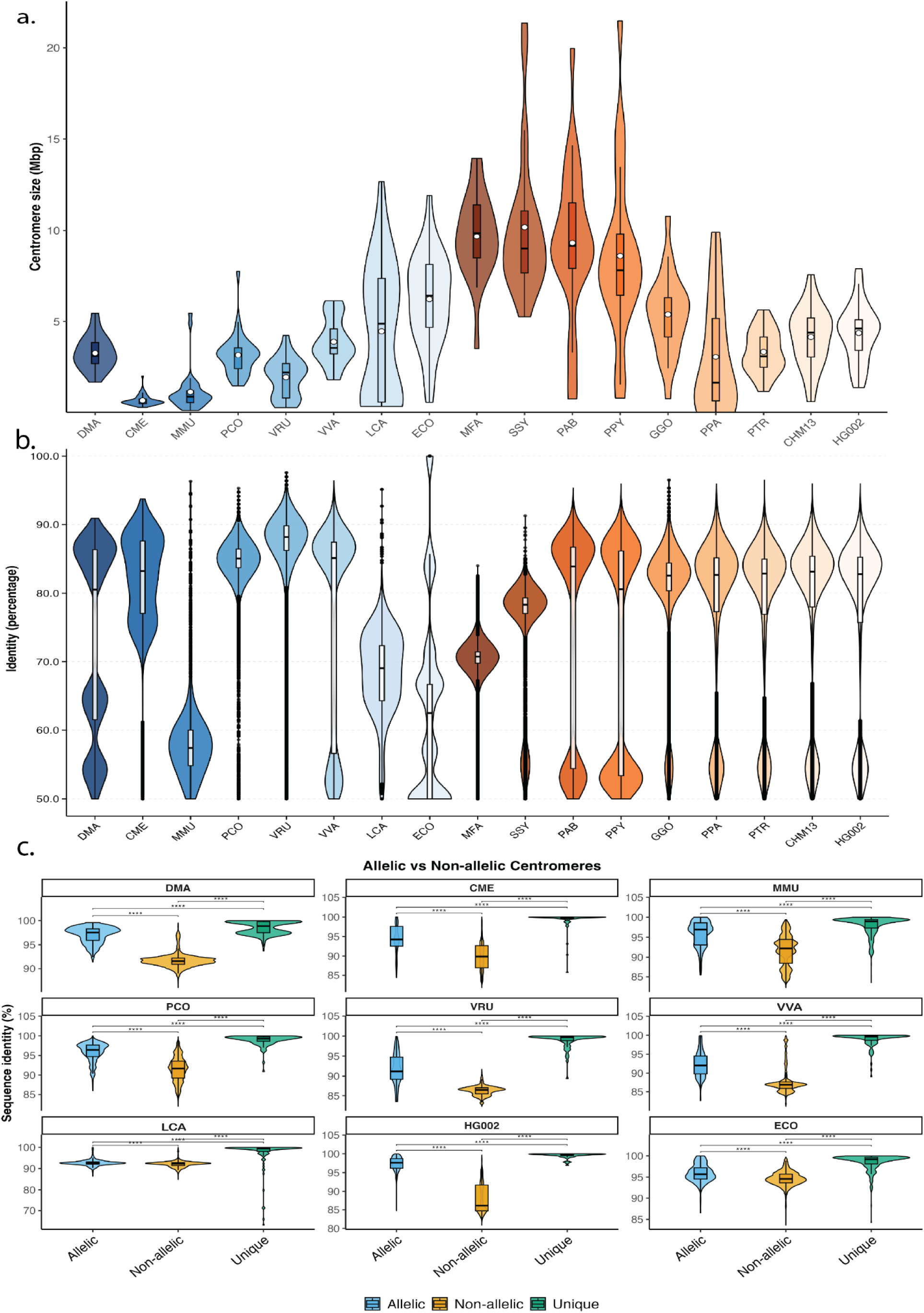
Comparative analyses of primate centromere length and divergence. a, Comparative centromere size distribution from lemurs, MFA and apes, including T2T-CHM13 and the diploid human genome HG002. b, Identity distribution of all the individual centromeric monomer units in a species, with respect to the most frequent monomer in that species. For non-lemur primates, canonical α-satellite was taken as the reference. c, Comparison between the identities of allelic and non-allelic centromeres in each species. Identities of unique sequences in every genome are plotted as control for the genome average.

**Figure 5.**
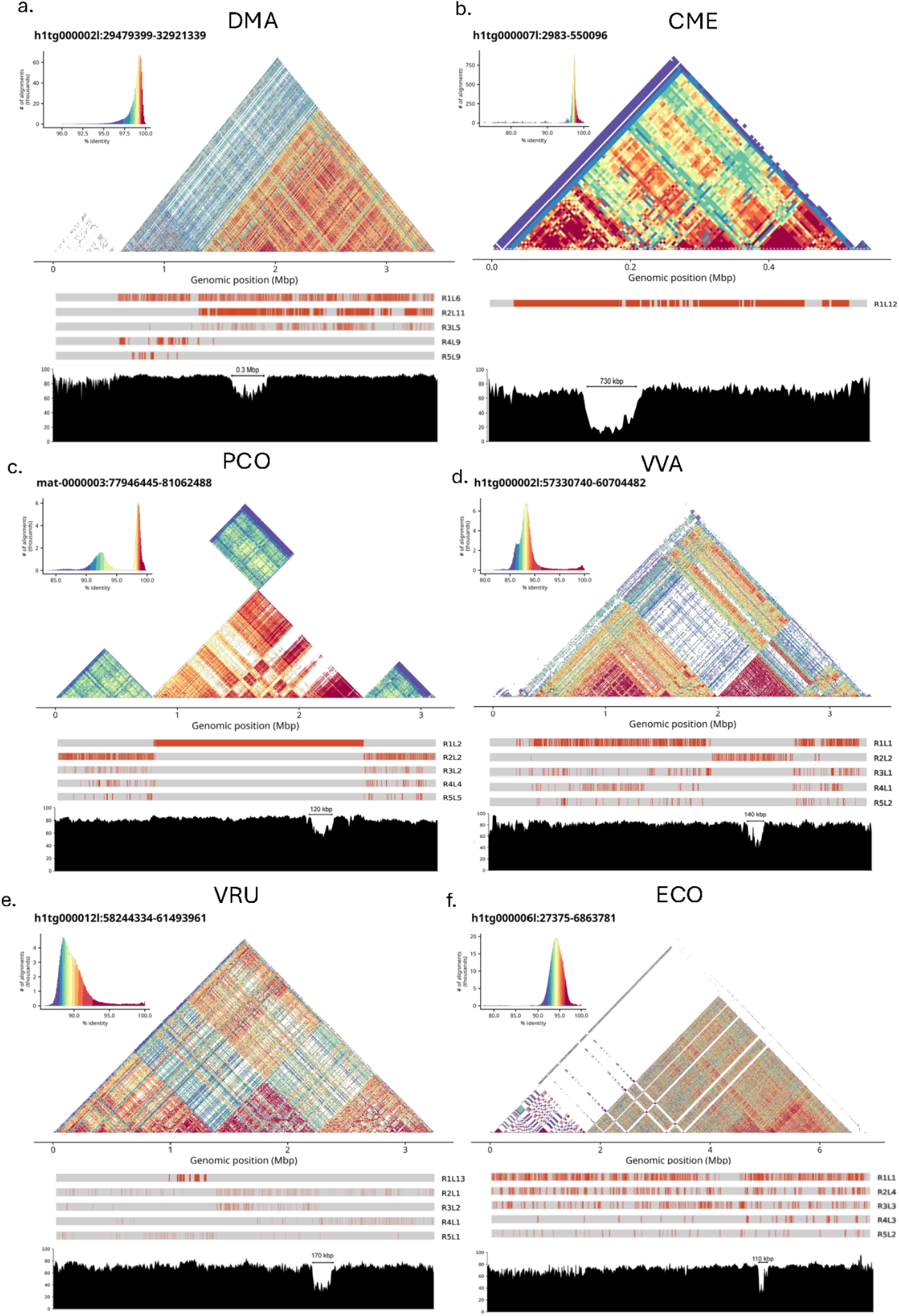
Lemur centromere higher-order repeat (HOR) structure and methylation. Heatmap of a centromere from each of six species (DMA, CME, PCO, VVA, VRU and ECO), along with their HOR annotations and 5-methylcytosine profiles. The HOR annotations show different HOR structures with score rankings as ‘R’ and length of individual HOR as ‘L’. CME has only a single HOR. Hypomethylation centromere dip regions and their estimated length (arrows) are shown in the context of the HOR structure and are variable between species.

**Figure 6.**
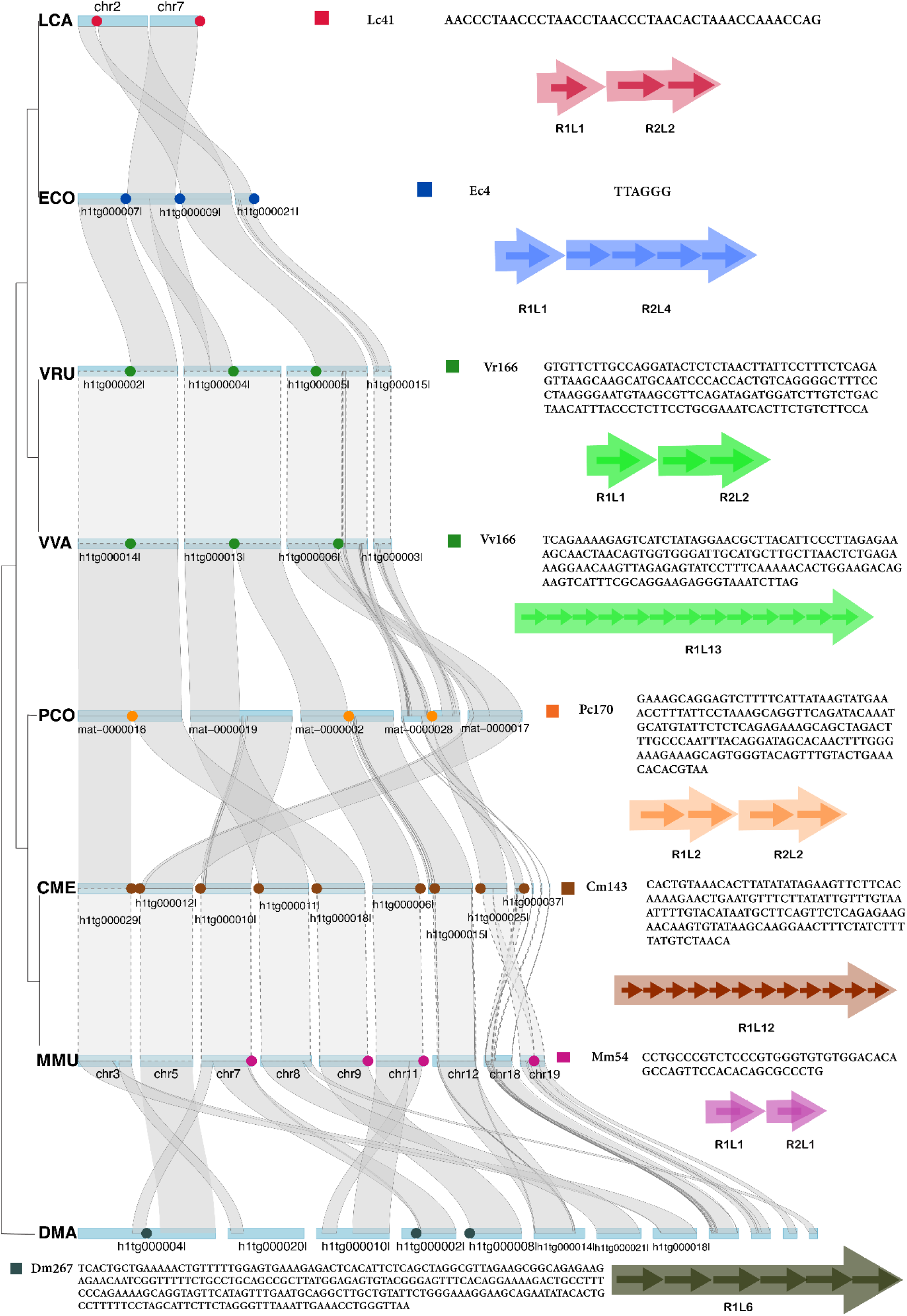
Centromere composition and repositioning with respect to synteny breakpoints among the lemurs. Mapping of two LCA chromosomes across all the related species showing the repositioning of centromeres and breakpoints of synteny across the lemur clade. Also described are the monomer sequences of each species and their most frequent HOR structure of each species. Dashed synteny lines across two species pairs demonstrate a near perfect synteny conserved between the phylogenetically closely related species.

**Table 2.** Summary of the assembled centromeres.

| Species | Centromere motif | Number of assembled centromeres (both haps) | Length | GC content | Most frequent HOR length |
| --- | --- | --- | --- | --- | --- |
| **Aye-aye (DMA)** | Dm267 | 19 | 267 | 43.82 | 6 |
|  | Dm209 |  | 209 | 40.67 |  |
|  | Dm222 |  | 222 | 39.64 |  |
| **Dwarf Lemur (CME)** | Cm143 | 64 | 143 | 28.67 | 3 |
| **Mouse lemur (MMU)** | Mm54 | 28 | 53 | 67.92 | - |
| **Coquerel's sifaka (PCO)** | Pc170 | 33 | 171 | 39.05 | 2 |
| **Ring-tailed lemur (LCA)** | Lc41 | 25 | 41 | 43.9 | - |
| **Red ruffed lemur (VRU)** | Vr166 | 20 (11 of only Vr166, 3 only Vr1405) | 166 | 42.17 | 1 |
|  | Vr1405 |  | 1405 | 46.05 |  |
| **Black and white ruffed lemur (VVA)** | Vv166 | 21 (16 of only Vv166, 3 of only Vv1399) | 166 | 42.17 | 1 |
|  | Vv1399 |  | 1399 | 45.46 |  |
| **Collared brown lemur (ECO)** | Ec6 | 32 | 6 | 50 | - |
|  | Ec548 |  | 548 | 47.08 |  |

Overall, our analysis shows that the basic strepsirrhine centromere motifs differ by 10-fold in length ranging from 41–548 bp (Table 2). Perhaps surprisingly, all species belonging to different genera harbor their own specific centromeric monomer sequence motifs. This suggests higher rates of centromere turnover than that observed for haplorhines. Both congeneric species belonging to *Varecia*, in contrast, share the same monomer repeat unit (Table 2).

In order to validate our candidates, we performed FISH (fluorescence in situ hybridization) and immunoFISH experiments for strepsirrhine species for which we had access to available cell lines for experimental testing (n=4). For example, we superimposed our centromere candidate sequences for *Lemur catta* (LCA) onto chromosomal scaffolds developed as part of the NCBI reference (mLemCat1, GCF_020740605.2). Similar to haplorhines, we observe a notable reduction in transposon/retrotransposon density corresponding to the long tandem repeat arrays (Figure 3). To confirm that these repeat arrays define centromeres, we constructed fluorescently labelled DNA probes specifically based on the LCA monomer for FISH imaging. Probes constructed with 41 bp monomers (repeat Lc41) were hybridized to metaphase spreads generated from LCA cell lines simultaneously labelled with CENP-C monoclonal antibodies. These immunoFISH experiments reveal that CENP-C antibodies and Lc41 FISH probes co-localize with the position of the primary constriction defining the centromeres. As expected, based on the telocentric lemur karyotype (Chu and Swomley, 1961), we find the centromere and Lc41 tandem repeat arrays located at the ends of chromosomes depicted in the karyotype (Figure 3a). Of interest, we failed to identify centromeres for four LCA chromosomes, including chromosome X. We presume that these chromosomes may have a minuscule centromere or potentially consist of non-repetitive neocentromeric DNA (Ventura et al., 2001; Cappelletti et al., 2025), and thus, would not have been detected by our approach.

We repeated this experimental validation approach for three additional lemur species (PCO, VVR and a *Eulemur* sp.) where cell lines could be obtained (courtesy of Christian Roos), performing co-immunoprecipitation experiments with CENPC antibodies and FISH DNA-probes designed to the predominant centromeric repeat identified from genome sequence and assembly. In all cases, centromeres or pericentromeric signals were confirmed (Figure 3b,c). Due to lack of an available cell line for *Eulemur collaris*, we used a closely related species instead, the gray-headed lemur (*Eulemur cinereiceps*), which confirms the dual role of telomeric repeats to define both the ends of the chromosome and the centromere associated with kinetochore binding (Figure 3d). See below for more detail.

### Comparative primate centromere and structure analyses

#### Aye-aye (DMA)

We assembled 19 centromeres in DMA, out of 30 total chromosomes. We distinguish three different but inter-related candidate centromeric repeats in DMA, whose sizes are 267 bp (Dm267), 222 bp (Dm222), and 209 bp (Dm209). We find that all the repeats have about 50% global pairwise sequence identity with each other but have a nearly identical core region of 75 bp, which aligns pairwise with greater than 95% identity in all three sequences. When compared to previously described monomers, DMA1 (146 bp) and DMA2 (268 bp) from Lee et al. (2011), DMA2 is identical to Dm267, but DMA1 makes partial sequences of all the other three sequences, along with DMA2 (Supplementary Figure S5a). We find that the centromeres consist of at least two of the three Dm sequences: Dm267 and Dm209, usually tandem repeats of one form and/or alternating with each other (see Supplementary Figure S5b and Supplementary Figure S4c for higher resolution views of the two centromeres). Overall, Dm267, Dm209 and Dm222 account for 43%, 32% and 14% base pair sequence, respectively, in all the complete centromeres. The other ∼11% of the centromere contains unique sequences. The most frequent monomer, Dm267, is organized into an R1L6 HOR structure in 15 out of 19 complete centromeres, with different secondary HORs in different centromeres. 5-methyl cytosine analysis shows distinguishable CDR for 17 assembled DMA centromeres, with the other two having small tentative dips. We estimate that CDRs for DMA range from 100 kbp to 300 kbp, with an average of 180 kbp (Figure 5a).

#### Dwarf lemur (CME)

For this species, we recovered the largest number of centromeres, with 64 of 66 chromosomes showing complete centromere assembly. The predominant centromere motif, Cm143, is a highly AT-rich (70.3%) monomer. The centromere structure consists of only one monomer, though of note, we find 11 centromeres that consist of two highly diverged HORs with identity of 72% between both the repeat arrays (Supplementary Figure S7). All the other chromosomes contain a single prominent HOR structure, with R1L3 being the most common (31 centromeres) followed by R1L12 in seven centromeres and another 26 centromeres having varied HOR configuration (as confirmed by CENdetectHOR; Daponte et al., 2025, Supplementary Table 5). The average size of CME centromeres at 520 kbp is significantly smaller than that observed for other lemur species (p < 0.01) (Figure 4a), but proportionally the CDR is larger. For example, the centromere in Figure 5b has a CDR 130 kbp long, which is about 38.4% of the total centromere size. Other 57 centromeres have a single CDR, six have two CDRs each, and one was observed with three CDRs.

#### Gray mouse lemur (MMU)

For this species, the monomer was previously described as Mm53, a 53 bp repeat (Larsen et al., 2017). We find a similar repeat consensus but with one additional base pair, thus relabelling it Mm54. We successfully characterize 28 centromeres from the whole karyotype of 66 chromosomes. There is no prominent HOR structure in MMU centromeres. We find CDRs in five centromeres only, with inconsistent methylation pattern for the remaining other 28 centromeres (Supplementary Figure S13). The basis for this reduced CDR signal for the mouse lemur is unknown. We note, however, that the DNA used for this particular sample was extracted more than 4 years ago and methylation signals are known to degrade over time (Lee et al., 2023).

#### Coquerel’s sifaka (PCO)

Sifaka has a 170 bp consensus centromere motif sequence (Pc170), similar in size to the α-satellite monomer in haplorhines, but there is otherwise no homology. We were able to assemble 33 centromeres out of 48 total chromosomes.The availability of parental data allowed this assembly to be fully phased and we successfully assigned 16 centromeres to the paternal haplotype and 17 to the maternal haplotype. All sequence-resolved centromeres in PCO show a well-defined dimer array: R1L2 is the predominant HOR and the same HOR structure was identified by CENdetectHOR (Daponte et al., 2025), with only three centromeres as exceptions (Supplementary Table 5). HORs are organized around a high sequence identity core, flanked by lower, more divergent flanking satellites. The CDR is very clearly demarcated in the centromeres, also overlapping the aforementioned high-identity region. All the CDRs are consistently between 110–150 kbp in all the centromeres (Figure 5c).

#### Black and white ruffed lemur (VVA)

Candidate assembled centromeres decompose into two monomers of distinct size: 166 bp (Vv166) and a much longer 1399 bp (Vv1399). From the expected karyotype of 46 chromosomes, we assembled 21 candidate centromeres, of which 16 consist solely of Vv166, three only of Vv1399, and the other two comprised of a fusion of both Vv166 and Vv1399 arrays in tandem, or one flanking the other (Supplementary Figure S8). For the centromeres consisting only of Vv166, R1L1 is the most frequent HOR configuration in six centromeres, but there was considerable variation in other HOR configurations found in other centromeres, ranging from R1L3 to R1L35. For the Vv1399 centromere, there is only one HOR, R1L1. While examining heterochromatin methylation, we observe an unexpected CDR depending on the type of centromeric arrays. We note a single canonical CDR for Vv166-based centromeres (Figure 5d), in contrast to multiple narrower CDRs for centromeres composed of Vv1399 or mixed/fused Vv1399/Vv166 (Supplementary Figure S9). This may suggest a sequence dependence on the pattern of CDR formation. Co-ImmunoFISH showed perfect colocalization between the probe Vv166 and CENP-C protein in 9 out of 24 chromosomes, while the probe Vv1399 displayed neither colocalization with immunohybridization signals nor with Vv166 FISH probes. Instead it identified pericentromeric loci for 5 out of 24 Vva chromosomes (Figure 3c). These results suggest a dual composition of *Varecia* centromeres: Vv166-associated functional centromeres, clearly marked by CENP-C colocalization, and centromeres where we were not able to identify satellite components in chromosomes lacking Vv166 signal. Vv1399, instead, is best interpreted as a pericentromeric satellite without detectable association with centromeric function. One possibility may be that the lack of detectable FISH signals for Vv166 for other chromosomes may result from the presence of satellite arrays whose size in these chromosomes is below the resolution of FISH.

#### Red ruffed lemur (VRU)

Given that this congeneric species diverged from VVA recently during the Pleistocene period (dos Reis et al., 2018), the predominant monomers are, unsurprisingly, homologous to VVA. Following the pattern of VVA, VRU centromeres consist of two monomers, which are nearly identical to VVA. For the longer one, the consensus motif has lengthened by 6 bp (Vr1405), while the other, more abundant motif remains the same in length (Vr166). We assembled 20 centromeres in VVA, where 11 are composed almost entirely of Vr166, three are composed of only Vr1405, and six contain both in which small arrays of Vr1405 are flanking the Vr166 array. R1L1 is the most frequent HOR in the Vr166 monomer with four centromeres, with substantial variation of HOR lengths in other centromeres. The methylation patterns also follow the same trend with Vr1405 centromeres having multiple CDRs in vicinity of each other, in contrast to a single CDR for the Vr166 centromeres (Figure 5e).

#### Collared brown lemur (ECO)

We discovered a particularly unexpected centromere composition in this species. Known telomere repeats of ‘TTAGGG’ form long interstitial arrays of average length 6.22 Mbp that define the putative assembled centromeres. This repeat (Ec6), along with sporadically interspersed Ec548 repeats, predominate in all 32 centromeres that we successfully assembled for ECO (2n ∼ 50). These interspersed filler regions are readily apparent in the StainedGlass heatmaps (Figure 5f). Due to the very small size of the repeat monomer, we could not distinguish HOR structures for centromeres except one that showed the R1L1 structure according to HiCAT, which implies that it does not contain any HOR structure. Nevertheless, we identify at least one CDR in every centromere, and for six centromeres we observe two CDRs. Co-immunoFISH experiments confirm that the telomeric repeat sequence has been co-opted to function as a centromere with typical kinetochore binding properties, while the satellite Ec548 appears to label pericentromeric satellite DNA, rather than as a sequence directly associated with centromeric activity (Figure 3d). The reuse of a telomeric repeat to define both the telomere and centromere within *Eulemur* may contribute to the high levels of variability in karyotype numbers seen in general within the brown lemur clade (Rumpler and Dutrillaux, 1976; Ventura et al., 2001).

#### Ring-tailed lemur (LCA)

For LCA, we scaffolded our assembly, using RagTag (Alonge et al., 2022), on the whole-karyotype assembly of 56 chromosomes from NCBI (GCF_020740605.2). We recovered 25 assembled centromeres where a 41 bp monomer (Lc41) predominates. We also identify other monomers extending the basic 41-mer to 76 bp and 117 bp. Due to this variation and interdigitation of non-canonical derivatives of Lc41 including smaller motifs, LCA centromeres show comparatively less sequence identity when compared to other LCA centromeres (Supplementary Figure S10). We did not find any prominent HOR structure in the LCA centromeres. Unlike most other lemur centromeres, we found no evidence of a clearly defined CDR for any centromere; instead, methylation patterns continuously fluctuate (Supplementary Figure S11). Co-immunoprecipitation experiments, however, confirm Lc41 as the centromere repeat motif (Figure 3a).

### Strepsirrhine CENP protein evolution

To assess whether centromere-associated proteins show signatures of adaptive evolution specific to lemurs, we performed clade-specific selection analyses on codon-aligned sequence for three genes encoding proteins critical for kinetochore function: CENP-A, CENP-B, and CENP-C (Methods). We partitioned the species tree into catarrhines, platyrrhines, and lemurs and treated each clade in turn as the “foreground.” Signals are strongest with CENP-B. With lemurs as “foreground,” PAML detects positive selection at approximately 9% of sites of CENP-B (ω = 3.00, p < 0.0001), against a backdrop of strong conservation across other primates (global ω = 0.10). Multiple independent tests corroborate this finding for CENP-B: RELAX indicates that positive selection strength on CENP-B is nearly twice as intense in lemurs when compared to other primates (K = 1.86, p = 0.0001) and BUSTED confirms gene-wide positive selection on lemur branches (p = 0.010). Importantly, neither catarrhines nor platyrrhines show comparable signals when tested as foreground, establishing that this is a lemur-specific phenomenon rather than a general primate pattern. Moreover, CENP-A shows site-specific positive selection in both lemurs (ω = 4.49, p = 0.003) and catarrhines (ω = 5.99, p = 0.021), with RELAX and BUSTED yielding non-significant signals. CENP-C shows a significant lemur signal in PAML and BUSTED (p = 0.001 and p = 0.005) but the RELAX result is not significant. Together, these results indicate strong positive selection on *CENPB* specific to lemurs, with weaker, less consistent signals for *CENPA* and *CENPC*.

## DISCUSSION

Centromeres harbor some of the most rapidly evolving repetitive sequences in eukaryotic genomes, and among primates most of their characterization has been restricted to the Old World Monkey lineages (Csink and Henikoff, 1998; Logsdon et al., 2024). We apply long-read sequencing methods to eight lemur species to investigate centromere evolution, repositioning, and chromosomal structural variation at an unprecedented resolution. The haplorhine lineage, comprising Platyrrhini, Catarrhini, and Tarsiidae, whose common ancestor lived ∼67-75 million years ago (dos Reis et al., 2018), is unified by α-satellite DNA as the predominant centromeric repeat, with variation observed among HOR organization between apes, Old World and New World monkeys (Yoo et al., 2025; Cacheux et al., 2018; Zhang et al., 2025; Nishihara et al., 2021). In contrast, lemurs exhibit a fundamentally different centromeric landscape: we identify seven distinct satellite monomer families across eight species spanning ∼41 million years, with no motif shared across distinct genera. This implies turnover of the predominant centromeric repeat at ∼1.7 motif changes per 10 million years, far exceeding haplorhine conservation (1 motif/∼40 million years). This is consistent with more rapid centromere evolution, potentially as a result of an ongoing centromere drive arms race. Whether centromeric repeats are a cause or consequence of centromere evolution remains debated, with evidence supporting a meiotic drive-mediated competition and feedback loop between satellite DNA and histone proteins, an evolutionary arms race underlying the centromere paradox (Henikoff et al., 2001; Cooper and Henikoff, 2004; Logsdon et al., 2020).

In our study, the convergence of a positive selection model on CENP-A and CENP-B in lemurs is consistent with the centromere drive model and provides the clearest molecular evidence yet for an ongoing arms race. Moreover, the signal for CENP-B emerges against a background of strong purifying selection, consistent with the broad mammalian conservation of CENP-B reported in previous studies (Sullivan and Glass, 1991; Okada et al., 2007), making its lemur-specific departure all the more compelling. This has direct relevance to the proposed role of centromere incompatibility in lemur speciation: if CENP-B was evolving adaptively in the ancestors of extant lemur clades, centromere recognition machinery was diverging concurrently with the karyotypic changes, providing a molecular mechanism through which centromere drive could have contributed to hybrid meiotic dysfunction and reproductive isolation.

We contend that this arms race, centromere repositioning, and overall chromosomal evolution have had a major role in the diversification of lemur species and may be consistent with White’s model of stasipatric speciation (White et al., 1968; Kearney and Hewitt, 2009). White championed the idea of stasipatric speciation in his writings, but it remained a fringe theory due to lack of data explaining fixation and spread of chromosomal variants (White, 1968; White, 1978; King, 1993). High species-specific centromere turnover rates from this study as well as previous observations of highly variable karyotype numbers and multiple fissions and fusions between closely related lemur species may all be linked (Rumpler and Albignac, 1975; Kolnicki, 2000; Rocchi et al., 2012). There is ample evidence that chromosomal rearrangements suppress recombination from studies in apes (Lin et al., 2025; Porubsky et al., 2020), deer mice (Hager et al., 2022), wild mice (Marin-Garcia et al., 2024), and muntjacs (Yin et al., 2021), consistent with White’s stasipatric model. Meiotic drive is considered to be the mechanistic basis of selecting favored states of karyotypes, mainly either predominantly acrocentric or metacentric (Pardo-Manuel de Villena and Sapienza, 2001; Blackmon et al., 2019). We hypothesize that meiotic drive may be a core mechanism in lemur speciation, ensuring that particular variants of centromeres get selected particularly during female gametogenesis and become fixed in the parent population along with the chromosomes that carry them. Meiosis in females is asymmetric and can select for the particular variants that favor the egg over the polar bodies. Later, these divergent centromere monomers in different populations would cause differential CENP protein loading and meiotic dysfunction in hybrids, resulting in reproductive isolation despite all incipient lemur species residing within the same geographic island.

We further suggest that this process is concomitant with formation of evolutionary new centromeres, a process previously indicated by FISH studies on *Lemur catta* and *Eulemur fulvus* X chromosomes (Ventura et al., 2004; Rocchi et al., 2012), which should also have accentuated lemur diversification. Lemur karyotypes provide direct empirical support for this model. They have a preponderance of acrocentric chromosomes (Rumpler and Dutrillaux, 1976; Horvath and Willard, 2007), and a putative role of Robertsonian translocations in lemur evolution has been suggested since their first karyotypic characterizations (Chu and Swomley, 1961). This is now confirmed at sequence resolution (Figure 6), where syntenic blocks have broken across all species at centromere junctions, providing evidence of ancestral inter-specific Robertsonian translocations and centromere repositioning. Across all seven genera, centromeres have shifted from their syntenically conserved positions, except in both *Varecia* species, where sequence and positional conservation are maintained, and in Cheirogaleidae, where only synteny is conserved—consistent with their comparatively lower rates of karyotypic diversification.

Evolutionary new centromere formation is most strikingly observed in the *Eulemur collaris* (ECO) genome, where telomeric repeats appear to be functioning as centromeric repeats, demonstrating recent chromosome rearrangement events. The *Eulemur* genus is known to have a varied karyotype as well as high rates of speciation (Ventura et al., 2001), and these results strengthen the case that telomeric repeats play a prominent role in centromere and karyotype evolution more broadly (Villasante et al., 2007). The presence of telomeric repeat sequences at both the centromeres and the ends of chromosomes may relate to the high levels of variability in karyotype numbers reported in general for the brown lemur clade (Rumpler and Dutrillaux, 1976; Ventura et al., 2001).

This is not to say that allopatric speciation also did not play a significant role in lemur diversification. Past geographical events, especially during the Oligocene and Miocene periods, likely contributed to separating populations (Antonelli et al., 2022). Additionally, the emergence and expansion of new ecological niches—including grasslands alongside dry and humid forests, with rivers acting as further barriers—may have promoted lemur speciation and the overall biodiversity of the island (Goodman and Ganzhorn, 2004; Igea and Tanentzap, 2021; Razafindratsima et al., 2025). We posit that, alongside these abiotic factors, molecular mechanisms such as centromere repositioning and chromosomal rearrangements provided speciation plasticity among phylogenetically related incipient species in close geographic proximity. The remarkable karyotypic diversity observed across lemur genera, and the species-specific centromere monomers and their rapid evolutionary turnover, are consistent with repeated shifts in the direction of meiotic drive, making lemurs an exceptional natural system in which to observe this process at genomic resolution. Linking chromosomal rearrangements and centromere evolution to macroevolutionary processes like speciation is a challenge, and empirical experiments with higher animals are simply not feasible (Harvey et al., 2019). These eight genome assemblies, however, open the door to functional studies of CENP binding divergence across species, hybrid fitness predictions from sequence data alone, and a comparative framework for testing centromere-driven speciation.

## ACKNOWLEDGEMENTS

We are thankful to Dr. Christian Roos at Deutsches Primatenzentrum, Göttingen, Germany, for his generosity in providing cell lines for FISH experiments. We thank the photographers of the lemurs at the Duke Lemur Center: Steve Coombs, David Haring, Sara Nicholson and Bob Karp. We also thank T. Brown for editing the manuscript and supplementary materials. This is Duke Lemur Center publication #XXXX. Research reported in this publication was supported, in part, by the Weill Neurohub Family Foundation and the National Human Genome Research Institute of the National Institutes of Health (NIH) under Award Number R01HG002385 (to E.E.Eichler). The content is solely the responsibility of the authors and does not necessarily represent the official views of the NIH. E.E.Eichler is an investigator of the Howard Hughes Medical Institute.

This article is subject to HHMI’s Immediate Access to Research policy, which requires that this article be made publicly available as initial and revised preprints deposited on a designated preprint server under a CC BY 4.0 license.

## DATA AND CODE AVAILABILITY

The raw genome sequencing data and assembly data generated for this project are available from Genbank with BioProject identifiers PRJNA1459402 - PRJNA1459415 and accession numbers SAMN58406792 - SAMN58406798. The gray mouse lemur (*Microcebus murinus*, MMU) genome is available under the accessions: GCA_040939455.2 and GCA_040939475.2. The assembly evaluation pipeline is available on (https://github.com/EichlerLab/assembly_eval) and (https://github.com/EichlerLab/assembly_qc). Code for centromere identification and selection analysis is available on (https://github.com/trihim/lemur_centromere).

## AUTHOR CONTRIBUTIONS

M.T., M.V., and E.E.Eichler conceptualized the study. E.E.Ehmke collected the samples. M.A., K.M.M., and K.H. generated the data. M.T., F.G., L.dG., D.Y., S.C., and C.L. conducted formal analyses. M.T., F.G., and L.dG. created visualizations. M.T., F.G., M.V., A.D.Y., and E.E.Eichler did the interpretation. M.T., F.G., M.V., and E.E.Eichler wrote the original draft. M.V. and E.E.Eichler supervised the study. M.K. and E.E.Eichler provided the resources. All authors reviewed and edited the manuscript.

## Supplementary Material

### Methods

#### A.#Genome Assembly

**1. Lemur DNA sequencing and genome assembly.** Blood samples were collected from seven lemur species (all except *M. murinus*) at the Duke Lemur Center (Duke IACUC protocols A208-23-10, A010-25-02; DLC protocol BSM-11-24-4) and used for DNA extraction. High molecular weight gDNA was extracted from frozen blood aliquots using the NEB Monarch HMW DNA extraction kit for Cells & Blood (#T3050L) following the manufacturer’s protocol. All individuals were sequenced using both PacBio HiFi long reads and Oxford Nanopore Technologies (ONT) ultra-long reads. Haplotype-resolved assemblies were generated using hifiasm v0.19.9 with HiFi and ONT reads for six species. For two species with available parental Illumina data (*P. coquereli* and *L. catta*), assemblies were independently generated with both hifiasm v0.19.9 and Verkko v2.2.1; Verkko assemblies were selected for downstream analyses on the basis of superior assembly contiguity and base-level accuracy. All assemblies were processed through a standardized quality-control pipeline (https://github.com/EichlerLab/assembly_qc). Assembly completeness and error profiles were evaluated using NucFreq (https://github.com/EichlerLab/assembly_eval) and Flagger v0.3.2 (Vollger et al., 2019; Liao et al., 2023). Collapsed regions were defined as loci where the second most frequent base exceeded 5× read depth, indicative of heterozygous sequence collapsed into a single haplotype. Duplicated and HiFi-depleted regions were defined as intervals with absent or markedly reduced HiFi read coverage, consistent with false duplication or assembly dropout, respectively. Assembly quality metrics for all eight species are provided in Supplementary Table S2. For *Lemur catta*, we used RagTag (Alonge et al., 2022) to scaffold the assembly on the reference genome in NCBI – GCA_020740605.1. The chromosomes in our assembly now follow the numbering from Cardone et al., 2002, which were mapped according to the synteny with human T2T-CHM13 genome.

A phased, haploid mouse lemur genome assembly was generated from primary fibroblasts of an adult index female (Chang, Lalgudi, Yoo, Trivedi et al., in preparation). Briefly, genomic DNA was extracted from the index and both parents. The index was sequenced using HiFi, ONT, and Hi-C Illumina platforms, while parental genomes were sequenced using Illumina short reads. A phased scaffold-level assembly was produced with hifiasm (v0.23.0) using trio-binning and ONT integration, followed by Hi-C scaffolding with HapHiC (v1.0.7) to generate a chromosome-level assembly. Assembly quality was evaluated with NucFreq and Flagger per above.

#### B.#Phylogenetic Reconstruction and Whole-Genome Alignment

A distance-based phylogeny was constructed using Mashtree with 100 bootstrap replicates. Sketches were generated with an assumed genome size of 2 Gbp and a minimum k-mer depth of zero to retain all k-mers, using the command:

`mashtree_bootstrap.pl --reps 100 --numcpus 12 --outmatrix --genomesize 2000000000 $(<fasta.fofn) -- --min-depth 0 > mashtree.tre`

Each lemur assembly was aligned pairwise against two reference genomes—*Macaca fascicularis* (MFA) and the human T2T assembly (T2T-CHM13)—using AnchorWave. Gene annotations for all species were derived from BUSCO gene models produced by compleasm (Huang and Li, 2023) as part of the assembly QC pipeline described above and used as anchor points for the AnchorWave alignment. Final alignments were produced using the anchorwave proali command, generating output in MAF format. MAF files were subsequently converted to PAF format using wgatools (Wei et al., 2025) for compatibility with downstream synteny and breakpoint analyses. Synteny breakpoints were identified using an in-house script. Synteny blocks were defined with a maximum intra-block gap in the target sequence of 100 kbp, a minimum query alignment length of 1 Mbp, and a minimum individual alignment length of 10 kbp.

#### C.#Centromere Identification and Characterization

##### Centromere monomer identification

Centromeric tandem repeat arrays were identified using TRF (Benson, 1999) applied to all eight assemblies. For a subset of species, results were independently corroborated using TRASH (Wlodzimierz et al., 2023), which performs an equivalent analysis and generates analogous summary outputs. This approach follows the framework first established by Melters et al. (2013) for mammalian centromeric repeat identification and has previously been applied in identification of mouse lemur centromeres (Larsen et al., 2017). For each species, repeat consensus length was plotted against total array size. Centromeric repeat arrays were identified by visual inspection of these plots, with centromeric candidates expected to produce peaks at a fundamental repeat unit length and its integer multiples, reflecting the hierarchical structure of satellite DNA. The fundamental repeat monomer for each species was defined as the lowest-order multiple identified from the TRF plots. All sequences of that unit length were extracted from the genome and the most frequently occurring sequence was designated the species monomer. Each monomer was then used as a BLAST query against the full genome assembly to precisely localize centromeric repeat arrays. Where multiple overlapping or adjacent arrays were detected, the longest contiguous array was designated the centromere locus. Monomer sequences were independently validated using MEME (Bailey et al., 2015), with the monomer length explicitly provided as a constraint and the BLAST-defined centromere array as input sequence; results were concordant across all species. These candidate repeat arrays were projected onto chromosomal ideograms to assess genomic distribution. Centromeric identity of candidate arrays was confirmed by fluorescence in situ hybridization (FISH) using probes derived from the identified repeats, co-stained with a CENP-C antibody.

##### FISH validation of centromeric repeats

###### Primer design and PCR amplification of centromeric monomers

To generate species-specific probes for the putative centromeric repeats of *Lemur catta* (LCA), *Propithecus coquereli* (PCO), *Varecia Variegata* (VVA) and *Eulemur collaris* (ECO), custom primers were designed based on their respective monomeric sequences to specifically amplify these regions. The primer sequences and the probes size are listed in Supplementary Table 4.

Contrastingly, we targeted the *Eulemur collaris* (ECO) 6 bp monomer, Ec6 (5’-TTAGGG-3’), using a pre-synthesized PNA-Cy3 probe from PNA Bio, Inc.

Genomic DNA was extracted from LCA, PCO, VVA, and ECI fibroblasts cultured in RPMI medium supplemented with 16% Fetal Bovine Serum (FBS), 1% L-Glutamine, and 1% Penicillin-Streptomycin. DNA was isolated using the QIAamp DNA Blood Mini Kit (Qiagen) according to the manufacturer’s protocol, quantified using a Nanodrop spectrophotometer (Thermo Fisher Scientific), and stored at –20°C.

PCR amplifications were performed using Thermo Scientific DreamTaq PCR Kit in 25 µL reactions containing 1× Master Mix, 0.25 µM of each primer, and 5 ng genomic DNA. For each reaction, thermal cycling included an initial denaturation at 95°C for 3 min, followed by 35 cycles of 95°C for 30 s, annealing at 55–60°C (depending on the species; see Supplementary Table 3) for 30 s, and 72°C for 1 min, with a final extension at 72°C for 7 min. PCR products were verified by electrophoresis on a 1% agarose gel, confirming the expected sizes.

###### Probe labeling and hybridization

Each PCR product was fluorescently labeled by a single-cycle PCR incorporating Cy3-dUTP, Fluorescein-dUTP, or Cy5-dUTP depending on the experiment. The reaction was performed in 25 µL containing 1× Taq buffer, 2 mM MgCl₂, 0.2 µM primers, 0.2 mM dNTPs, 0.1 mM labeled-dUTP, 0.4% BSA, and 1.25 U recombinant Taq polymerase (recombinant) (Thermo Fisher Scientific, Molecular Biology Grade), using 1 µL of PCR product as template. For each reaction, thermal cycling included an initial denaturation at 94°C for 30 sec, followed by 1 cycle of 94°C for 30 s, annealing at 55–60°C (depending on the species; see Supplementary Table 3) for 30 s, and 72°C for 1 min, with a final extension at 72°C for 10 min.

FISH experiments were performed on metaphase spreads of LCA, PCO, VVA, and ECI. Each cell line was arrested in metaphase with colcemid, incubated in hypotonic KCl (0.56%) for 30 min, pre-fixed with methanol: acetic acid (3:1), centrifuged, and fixed again in methanol: acetic acid (3:1).

Chromosome spreads were dropped onto glass slides, aged at 90°C for 1.5 h and treated with 0.005% pepsin in 0.01 M HCl, followed by three washes, 5 minutes each, in 1× PBS, 0.5 M MgCl₂, 4% paraformaldehyde, and a cold-ethanol series (70%, 90%, and 100%).

Monomer-based probes were fluorescently labeled according to the target species:

- For *Lemur catta* (LCA), the Lc41 monomer probe was labeled with Cy3.
- For *Varecia variegata* (VVA), the Vv166 monomer and the Vv1405 probes were labeled with Cy3 and Fluorescein (Fx), respectively.
- In *Propithecus coquereli* (PCO), the Pc170 probe was labeled with Cy3 and the probe targeting the Y centromere with Fx.
- For *Eulemur cinereiceps* (ECI), the Ec6 monomer was targeted using the PNA-Cy3 probe, while the Ec548 monomer-base probe with Fx.

All probes were subsequently precipitated by ion-exchange alcohol precipitation without Cot DNA (being probes targeting repetitive regions). Pellets were air-dried and resuspended in a hybridization buffer (50% formamide, 10% dextran sulfate, 2X saline sodium citrate (SSC)). For single-probe FISH, each probe was hybridized independently. For co-hybridization experiments, the two probes were co-precipitated and applied simultaneously to the same metaphase preparations.

For most species, hybridization was carried out overnight at 37°C after denaturation for 2 min at 70°C in HYBrite^TM^ Vysis. The only exception was *Eulemur cinereiceps*; due to the requirements of the PNA probe, a modified protocol was applied with denaturation performed at 75°C for 10 min. Post-hybridization washes were performed at 60°C in 0.1× SSC (three times, 5 min each), with the exception of ECI where washes were carried out at 60°C in 2x SSC, 0.1% Tween (two times, 5 min each). At the end, the slides were stained with DAPI and covered with coverslip, and signals were detected with specific filters using a Leica DMRXA epifluorescence microscope equipped with a cooled CCD camera (Princeton Instruments). Finally, images were processed using Adobe Photoshop™ software.

###### Immuno-FISH with Cenp-C antibody

LCA, PCO, VVA, ECI cell lines were colcemid-treated and incubated in hypotonic KCl (0.56%) for 30 min, pre-fixed with methanol:acetic acid (3:1), centrifuged, and fixed in methanol:acetic acid (3:1). Chromosome spreads were dropped onto glass slides and incubated for 4 days at 37°C.

Slides were rehydrated in 1× PBS-azide buffer (10 mM NaPO4, pH 7.4, 0.15 M NaCl, 1 mM EGTA, and 0.01% NaN3) for 15 min at RT and washed three times in 1× TWEEN buffer containing 0.5% Triton X-100 and 0.1% BSA.

Mouse anti-CENP-C (Abcam, ab50974) monoclonal antibodies were diluted to 0.001 µg/µL and incubated on the slides (100 µL/slide) for 2 h at 37°C. Slides were washed three times in 1x KB buffer (10 mM Tris-HCl, pH 7.7, 0.15 M NaCl, and 0.1% BSA) and incubated with 100 µL/slide goat anti-mouse IgG secondary antibody conjugated to fluorescein (Fx) (Abcam, ab6785; 1:100 dilution) for 45 min at 37°C in the dark. After incubation with the secondary antibody, slides were washed three times with 1× KB for 2, 5, and 3 min, prefixed in 4% paraformaldehyde in 1× KB for 45 min at RT, and then fixed in methanol: acetic acid (3:1) for 15 min.

For each cell line, FISH was then performed using the species-specific monomer-based probes differently labeled.

For single-probe FISH, each probe was hybridized independently. For co-hybridization experiments, the two probes were co-precipitated, as previously described, and applied simultaneously to the same metaphase preparations after resuspension in hybridization buffer (50% formamide, 10% dextran sulfate, 2× SSC). Hybridization was carried out overnight at 37°C after denaturation for 8 min at 70°C in HYBrite^TM^ Vysis. Post-hybridization washes were performed at 60°C in 0.1× SSC (three times for 5 min, high stringency). Slides were counterstained with DAPI and imaged with a Leica DMRXA2 epifluorescence microscope. DAPI, Cy3, fluorescein, and Cy5 signals were acquired separately with dedicated filters, recorded as grayscale images, pseudocolored, and merged using Adobe Photoshop™.

###### Methylation and higher-order repeat (HOR) organization

ONT reads for all eight species were basecalled with Guppy 6.5.7, Dorado v1.0.2, or higher with CpG methylation calling enabled. We aligned all the ONT reads to assembled genome and then methylation profiles across centromere arrays were visualized to identify centromere dip regions (CDRs), defined as localized hypomethylated intervals within otherwise highly methylated satellite arrays that mark the site of kinetochore assembly. Profiles were plotted using CDR-finder (Mastrorosa et al., 2024) and CDRs, where present, were manually identified and annotated.

HOR structure was characterized for all assembled centromeres using HiCAT (Gao et al., 2023). HiCAT defines HOR units in the format R*n*L*m*: where the rank (R) is derived from a composite score that integrates centromere coverage with a penalty for over-compressed local nesting, ensuring that the highest-ranked HOR optimizes the balance between total coverage and repeat fidelity; the parameter L denotes the HOR length, defined as the total number of constituent monomeric units. HOR structure was successfully resolved for most centromeres across all eight species. Exceptions were *L. catta* (LCA) and *E. collaris* (ECO), for which HiCAT was applied to a single representative centromere per species due to the disproportionate size of their centromere arrays relative to their short monomer units (41 bp and 6 bp, respectively), which precluded full-genome HOR decomposition.

HORs were also identified using CENdetectHOR (Daponte et al., 2025). CENdetectHOR defines the resulting HORs using the nomenclature format, CnHn, where Cn represents the contig identification number (substituting the chromosome identifier typically used in anchored assemblies) and Hn indicates the length of the identified HOR unit. The analysis was conducted only on the two species, *Propithecus coquereli* and *Cheroilogaleus medius*, utilizing scaffold-level assemblies. A consensus sequence for the respective monomeric repeats was used as a reference. The analysis followed the standard detection protocol, using default parameters.

Visual and structural analysis of the outputs was performed using the PhyloTreeGUI graphical interface, which allowed for the reconstruction and inspection of HOR-based phylogenetic trees. The downstream analysis involved the systematic pruning of HOR trees to extract biologically significant structures. To ensure a balanced comparison, branches were selected at a consistent hierarchical level across the different trees. The selection process was guided by a coverage threshold, maintaining approximately 95% coverage of the repetitive regions.

For each assembled centromere, array size and sequence identity relative to the species-specific monomer were calculated. As a reference for comparative context with haplorhine centromeres, the canonical alpha satellite monomer sequence was used:

AATCTGCAAGTGGATATTTGGACCGCTTTGAGGCCTTCGTTGGAAACGGGAATATCTTCATATAAAAACTAGACAGAAGCATTCTCAGAAACTTCTTTGTGATGTGTGCATTCAACTCACAGAGTTGAACCTTCCTTTTCATAGAGCAGTTTTGAAACACTCTTTTTGTAG

As all assemblies were fully phased or pseudo haplotype-resolved, allelic centromere pairs could be identified for each chromosome. Pairwise sequence identity was calculated between allelic centromeres (intra-individual) and between non-allelic centromeres (inter-individual), with unique single-copy genomic sequences used as a specificity control. Centromere sequences were aligned using minimap2 with the following command, according to Logsdon et al. (2024):

`minimap2 -I 15G -K 8G -t {threads} -ax asm20 --secondary=no --eqx -s 2500 {ref.fasta} {query.fasta}`

Alignment identity statistics were extracted from the resulting PAF files using rustybam (https://github.com/vollgerlab/rustybam).

Centromere arrays were extracted and visualized as pairwise sequence identity heatmaps using StainedGlass (Vollger et al., 2022). Window size was selected for each species proportionally to the species-specific monomer length to ensure biologically meaningful resolution of repeat structure.

#### D.#Molecular Evolution of Centromere Proteins

Orthologs of CENP-A, CENP-B, and CENP-C were identified in each lemur assembly using miniprot, with *Lemur catta* reference protein sequences as queries (GCF_020740605.2). Coding sequences were extracted from the resulting alignments using gffread (-x flag) to obtain coding sequence (CDS) in the correct reading frame. Sequences were inspected for premature stop codons and frameshifts prior to alignment. We also procured CENP CDS for 15 other genome assembled species in NCBI: *Carlito syrichta* (GCF_000164805.1), *Rhinopithecus bieti* (GCF_001698545.2), *Papio anubis* (GCF_008728515.1), *Trachypithecus francoisi* (GCF_009764315.1), *Pongo abelii* (GCF_028885655.2), *Pan paniscus* (GCF_029289425.2), *Macaca fascicularis* (GCF_037993035.2), *Macaca mulatta* (GCF_049350105.2), *Chlorocebus sabaeus* (GCF_047675955.1), *Cebus imitator* (GCF_001604975.1), *Sapajus apella* (GCF_009761245.1), *Aotus nancymaae* (GCF_030222135.1), *Saimiri boliviensis* (GCF_048565385.1), *Callithix jacchus* (GCF_049354715.1), and *Homo sapiens* (GCF_009914755.1).

CDS were translated using *seqkit translate* (Shen et al., 2024), selecting the longest isoform in case of multiple isoforms with AGAT’s agat_sp_keep_logest_isoform.pl script (Dainat et al., 2021), aligning the protein sequences using MAFFT (Katoh et al., 2019), and the resulting amino acid alignment was used to guide codon-aware back-translation of the corresponding nucleotide sequences using pal2nal (Suyama et al., 2006). The final codon alignments were used as input for all downstream selection analyses. All selection analyses were conducted on the species tree derived from Timetree5 (Kumar et al., 2022).

##### PAML codeml

Selection analyses were performed using codeml from the PAML package (Yang, 2007). A model M0 (one-ratio) analysis was first run for each gene to estimate the genome-wide background dN/dS (ω) ratio across all branches. Branch-site model analyses were then conducted to test for episodic positive selection along foreground (lemur) branches. The branch-site test compares a model allowing ω > 1 on foreground branches (Model A) against a null model with ω constrained to 1, using a likelihood ratio test (LRT). Internal node labels were specified using the ete3 tree-manipulation framework to ensure correct foreground branch designation in the PAML control file. P-values were corrected for multiple testing using the Benjamini-Hochberg procedure.

##### RELAX

Relaxation or intensification of selection pressure on lemur lineages was assessed using RELAX (Wertheim et al., 2015) as implemented on the Datamonkey web server. RELAX fits a model with a selection intensity parameter K to foreground branches relative to reference branches, where K > 1 indicates intensified selection and K < 1 indicates relaxation. The root node was explicitly labelled as the reference branch ({Reference}) as required by RELAX. Statistical significance was assessed by LRT against a null model with K constrained to 1.

##### BUSTED

Evidence for episodic diversifying selection anywhere in the lemur foreground branches was tested using BUSTED (Murrell et al., 2015) as implemented on the Datamonkey web server. BUSTED tests whether at least one branch and site combination in the foreground has experienced positive selection (ω > 1), without requiring that selection be pervasive. Results were considered significant at p < 0.05.

## Supplementary Tables

**Table 3.** Tests of selection on CENP genes.

| Gene | foreground | M0 $\omega$ | LRT M1aM2a Significance | LRTM7M8 Significance | BS Significance | BS (fg) $\omega$ | BS proportion |
| --- | --- | --- | --- | --- | --- | --- | --- |
| **CENPA** | Lemur | 0.5003 | * | ** | ** | 4.4929 | 0.0445 |
| **CENPA** | OWM | 0.5003 | * | ** | * | 5.9905 | 0.03 |
| **CENPA** | NWM | 0.5003 | * | ** | ns | 2.5081 | 0 |
| **CENPB** | Lemur | 0.1041 | ns | *** | *** | 3.0049 | 0.0886 |
| **CENPB** | OWM | 0.1041 | ns | *** | ns | 11.3848 | 0.0014 |
| **CENPB** | NWM | 0.1041 | ns | *** | ns | 1 | 0 |
| **CENPC** | Lemur | 0.7057 | *** | *** | ** | 2.8099 | 0.0364 |
| **CENPC** | OWM | 0.7057 | *** | *** | ns | 1.6704 | 0.044 |
| **CENPC** | NWM | 0.7057 | *** | *** | ns | 4.8556 | 0.0055 |
| **TRIM5** | Lemur | 1.1519 | *** | *** | *** | 6.1327 | 0.0364 |
| **TRIM5** | OWM | 1.1519 | *** | *** | *** | 8.9436 | 0.0177 |
| **TRIM5** | NWM | 1.1519 | *** | *** | *** | 8.2967 | 0.0312 |
| **LYZ** | Lemur | 0.5877 | *** | *** | *** | 8.4768 | 0.0245 |
| **LYZ** | OWM | 0.5877 | *** | *** | ns | 2.6782 | 0.0999 |
| **LYZ** | NWM | 0.5877 | *** | *** | ns | 4.5516 | 0.0197 |

( $\omega$ - dN/dS ratio, LRT - likelihood ratio test, BS - branchsite, ns - not significant, \* $p < 0.05$ , \*\* $p < 0.01$ , \*\*\* $p < 0.001$ )

**Supplementary Table 1.** Assemblies of Lemuriformes in NCBI.

| **Species** | **Assembly ID** | **# of contigs** | **Contig N50** | **Assembly level** | **Genome coverage** | **Reference** |
| --- | --- | --- | --- | --- | --- | --- |
| ***Lemur catta*** | GCF_020740605.2 | 394 | 32.5 Mbp | Chromosome | 20.81x | Palmada-Flores et al., 2022 |
| ***Varecia variegata*** | GCA_028533085.1 | 1,029,863 | 40.2 kbp | Chromosome | 74x | DNA Zoo (unpublished) |
| ***Microcebus murinus*** | GCF_000165445.2 | 50,982 | 210.7 kbp | Chromosome | 221.6x | Baylor College (Unpublished) |
| ***Varecia rubra*** | GCA_963573675.1 | 112,503 | 37.4 kbp | Scaffold | 35x | Kuderna et al., 2023 |
| ***Eulemur collaris* (referenced as *Eulemur fulvus collaris*)** | GCA_963575015.1 | 111,525 | 39 kbp | Scaffold | 35x | Kuderna et al., 2023 |
| ***Propithecus coquereli*** | GCF_000956105.1 | 299,069 | 28.1 kbp | Scaffold | 104.7x | Baylor College (Unpublished) |
| ***Cheirogaleus medius*** | GCA_008086735.1 | 130,903 | 34.9 kbp | Scaffold | 110x | Williams et al., 2020 |
| ***Daubentonia madagascariensis*** | GCA_044048945.1 | 930 | 80.4 Mbp | Scaffold | 135x | Versoza and Pfeifer, 2024 |

**Supplementary Table 2.** NucFreq and Flagger statistics of the new assemblies.

|  | *Microcebus murinus* | *Cheirogaleus medius* | *Propithecus coquereli* | *Daubentonia madagascariensis* | *Lemur catta* | *Varecia rubra* | *Varecia variegata* | *Eulemur collaris* |
| --- | --- | --- | --- | --- | --- | --- | --- | --- |
| **NucFreq** |  |  |  |  |  |  |  |  |
| Misassembly | 3.08 Mbp | 0.75 Mbp | 1.33 Mbp | 0.65 Mbp | 0.37 Mbp | 0.35 Mbp | 0.18 Mbp | 0.78 Mbp |
| Collapse | 4.16 Mbp | 0.92 Mbp | 23.2 Mbp | 0.66 Mbp | 0.78 Mbp | 0.20 Mbp | 0.14 Mbp | 0.19 Mbp |
| **Flagger_0.3.3** |  |  |  |  |  |  |  |  |
| Erroneous | 13.2 Mbp | 2.86 Mbp | 17.3 Mbp | 7.09 Mbp | 7.76 Mbp | 1.57 Mbp | 1.25 Mbp | 8.25 Mbp |
| Duplicated | 140 Mbp | 93.1 Mbp | 18.7 Mbp | 43.1 Mbp | 48.5 Mbp | 34 Mbp | 53.5 Mbp | 141 Mbp |
| Collapses | 9.92 Mbp | 91.4 Mbp | 26.3 Mbp | 1.30 Mbp | 15.8 Mbp | 0.77 Mbp | 0.64 Mbp | 137 Mbp |
| Unknown | 1.22 Mbp | 0.32 Mbp | 0.21 Mbp | 0.94 Mbp | 0.25 Mbp | 0.09 Mbp | 0.12 Mbp | 0.30 Mbp |
| Haploid | 4.57 Gbp | 4.37 Gbp | 4.60 Gbp | 4.89 Gbp | 4.49 Gbp | 4.34 Gbp | 4.35 Gbp | 4.19 Gbp |

**Supplementary Table 3.** Mapping between chromosome numbers of Lemur catta karyotype from Cardone et al., 2002 and NCBI assembly.

| Cardone et al., 2002 | Lemur_NCBI |
| --- | --- |
| 1 | NC_059128.1 |
| 2 | NC_059129.1 |
| 3 | NC_059130.1 |
| 4 | NC_059132.1 |
| 5 | NC_059131.1 |
| 6 | NC_059139.1 |
| 7 | NC_059135.1 |
| 8 | NC_059134.1 |
| 9 | NC_059138.1 |
| 10 | NC_059133.1 |
| 11 | NC_059136.1 |
| 12 | NC_059141.1 |
| 13 | NC_059137.1 |
| 14 | NC_059140.1 |
| 15 | NC_059144.1 |
| 16 | NC_059147.1 |
| 17 | NC_059142.1 |
| 18 | NC_059143.1 |
| 19 | NC_059146.1 |
| 20 | NC_059145.1 |
| 21 | NC_059148.1 |
| 22* | NC_059150.1 |
| 23* | NC_059151.1 |
| 24* | NC_059152.1 |
| 25 | NC_059149.1 |
| 26* | NC_059153.1 |
| 27* | NC_059154.1 |
| X | NC_059155.1 |
| Y | NC_059156.1 |

**Supplementary Table 4.**
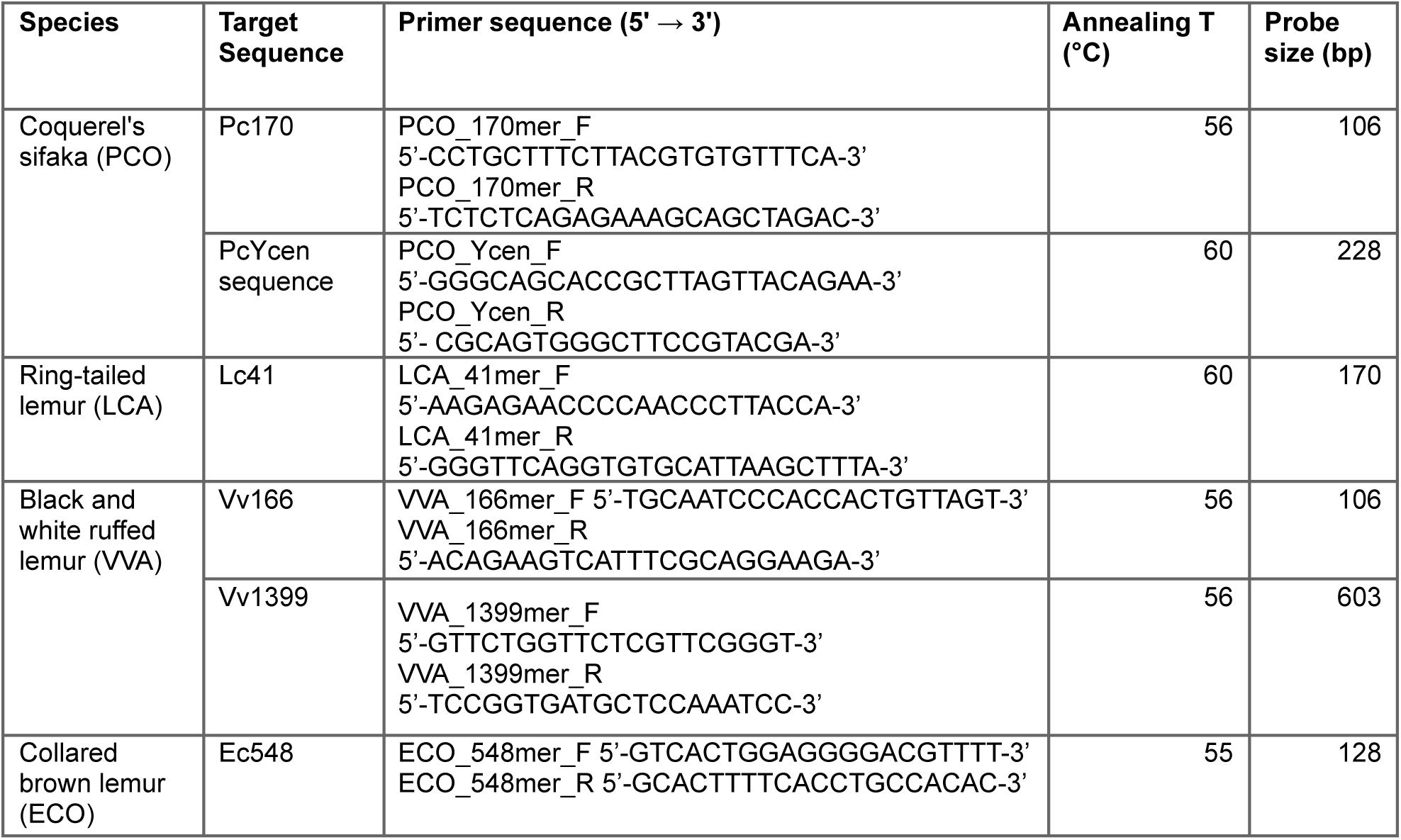
Primers used for probe synthesis. Annealing temperatures and expected product sizes are provided for each species-specific monomer targeting putative centromeric regions.

**Supplementary Table 5.**
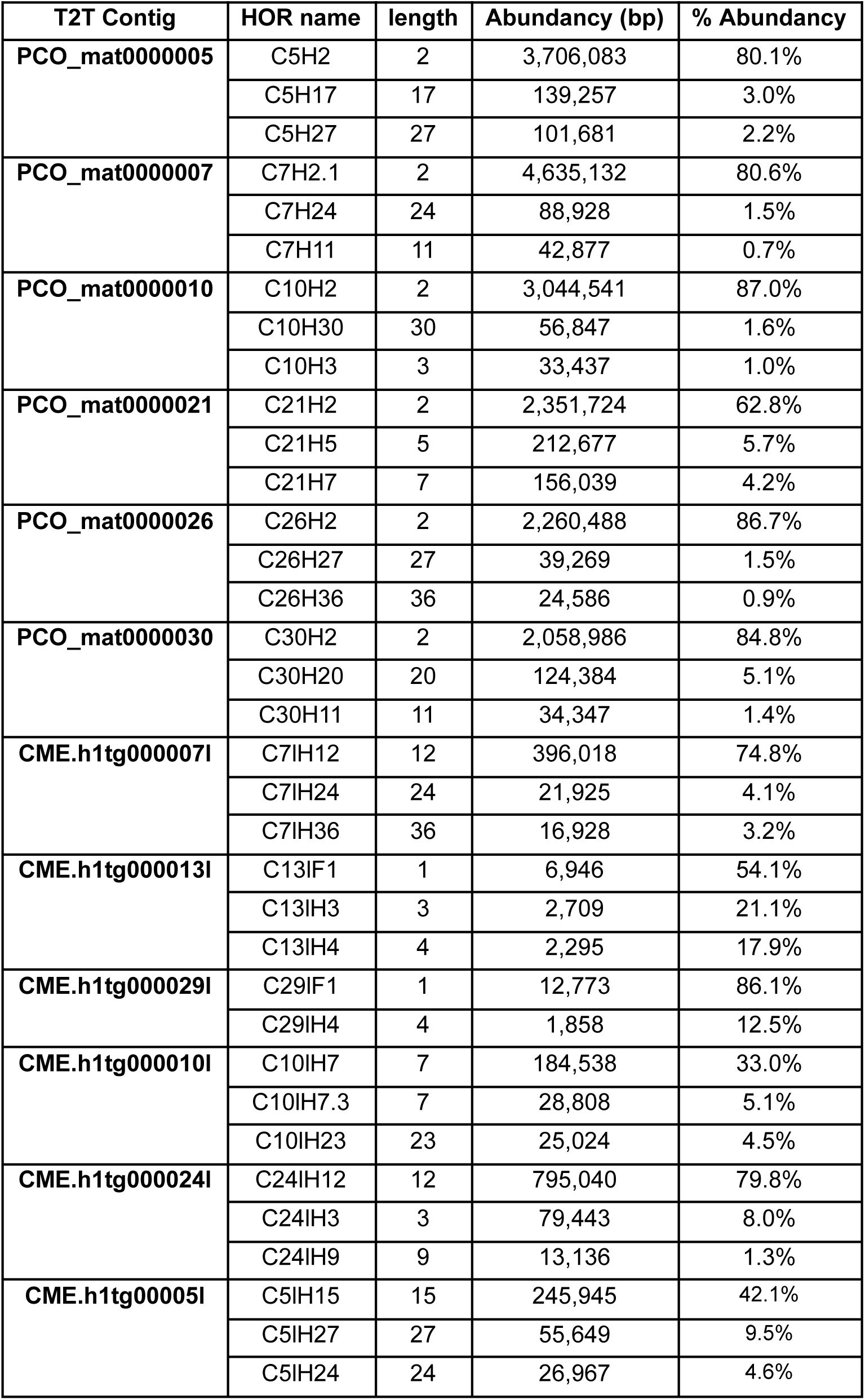
CENdetectHOR results for the T2T contigs of two species: CME and PCO.

## Supplementary Figures

**Figure S1.**
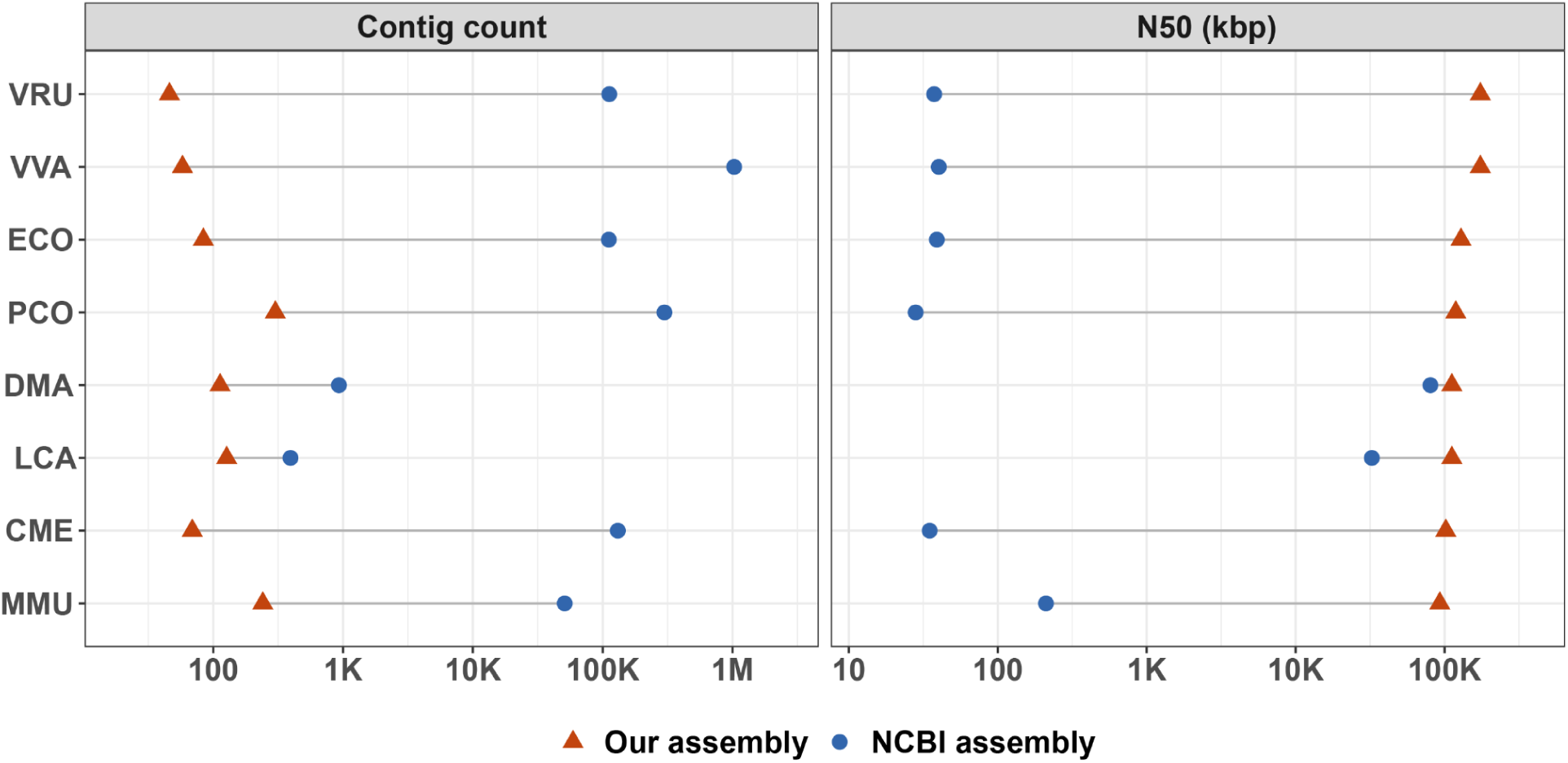
Improvement in assembly contiguity in this study. Comparison between the assemblies already submitted in the NCBI database for these lemur species, and the assemblies presented in this study. It can be clearly seen that both contig count and contig N50 sizes have drastically improved in our assemblies.

**Figure S2.**
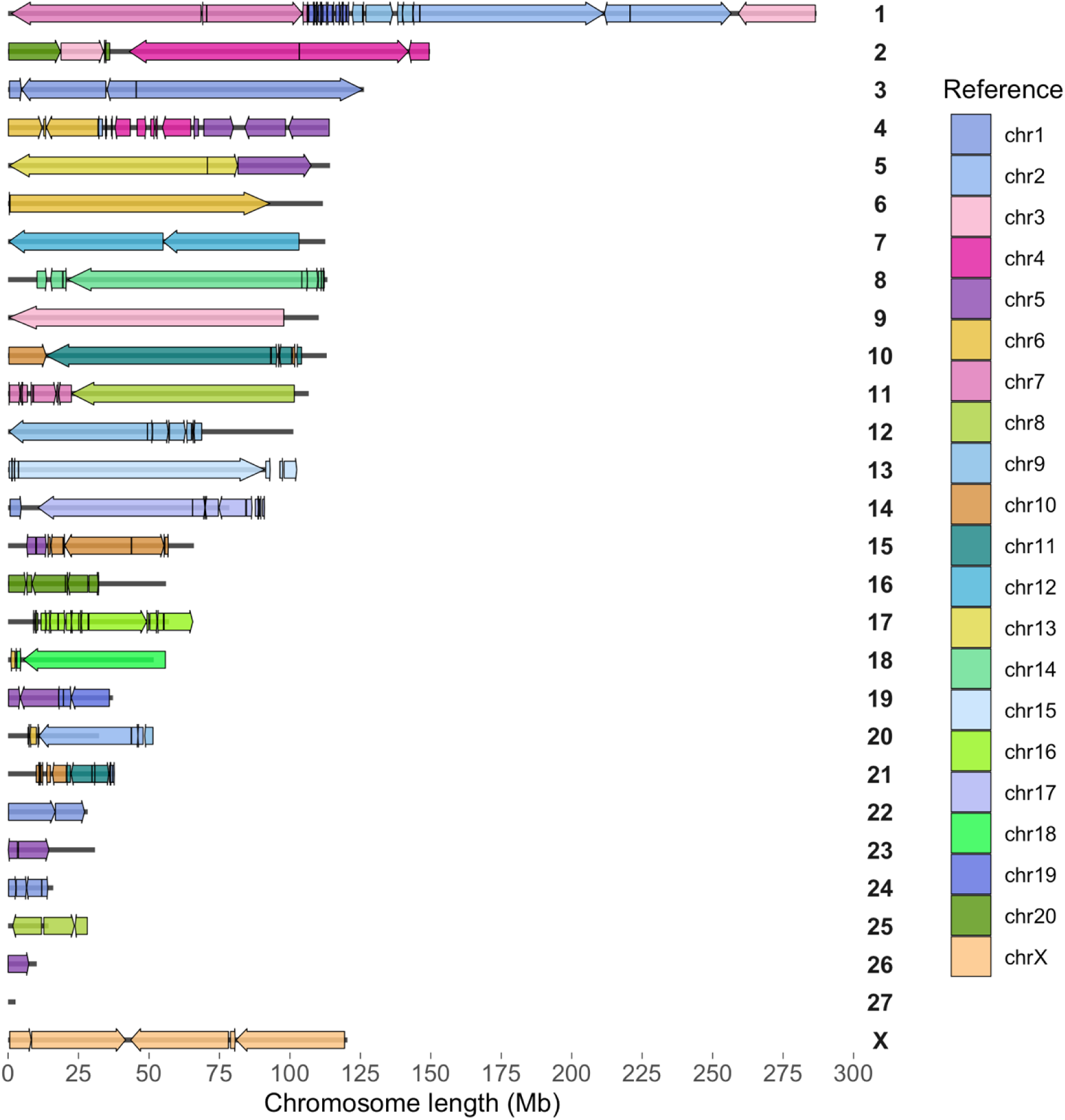
Synteny of. *Lemur catta* **genome with Macaca fascicularis (MFA).** Synteny between MFA and *Lemur catta* (LCA) chromosomes, with LCA on the left and MFA chromosome colors on the right. Arrows show the orientation of MFA synteny blocks and straight lines showing the regions with no syntenic matches. There are six chromosomes which are fully conserved in both the species, viz., 7 (chr14 in MFA), 8 (chr12), 10 (chr15), 15 (chr16), 20 (chr20) and X. Even within these chromosomes rearrangements are visible in both p and q arms with centromere position.

**Figure S3.**
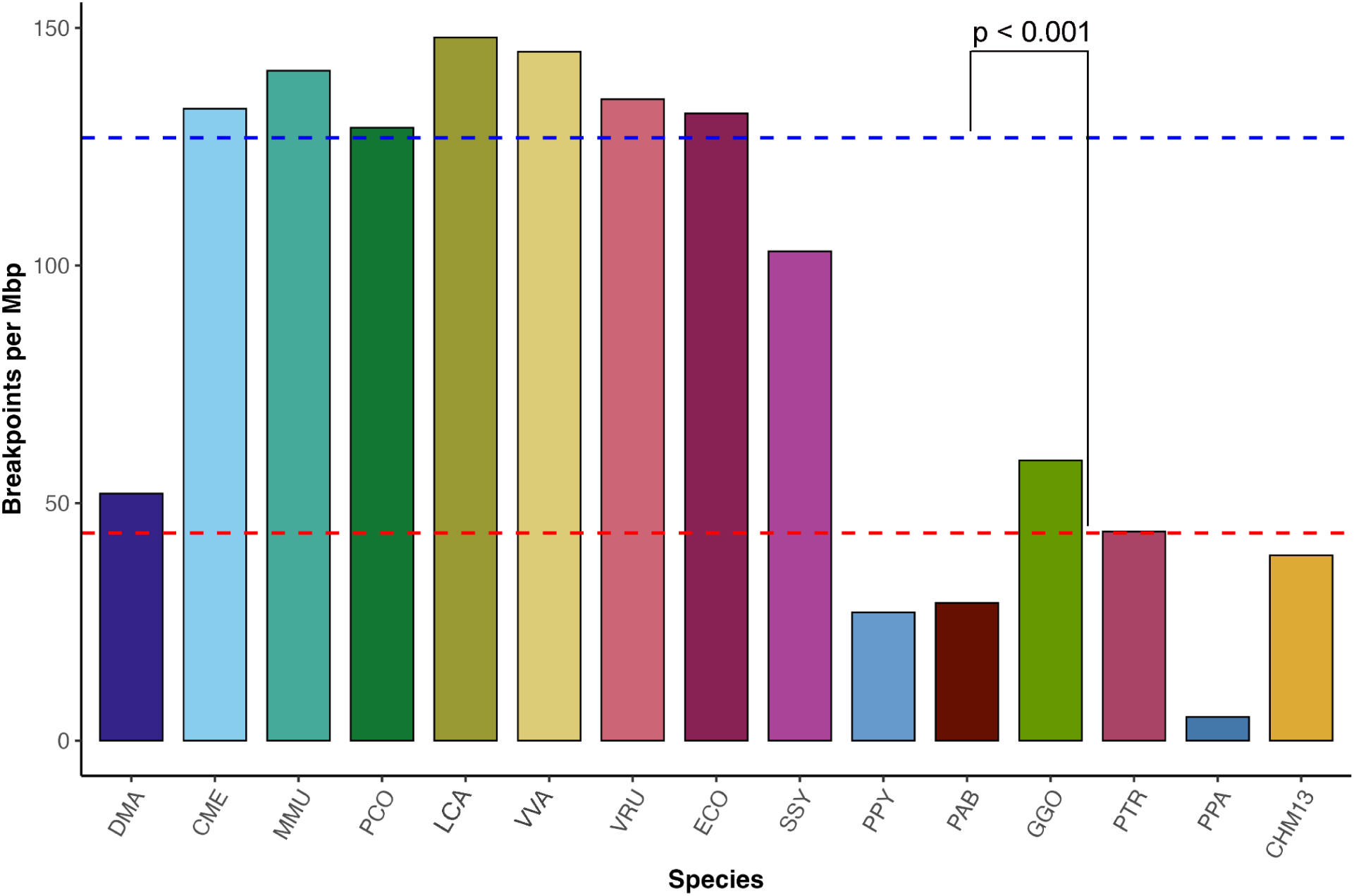
Breakpoints of lemur genome with respect to MFA. The plot displays the total breakpoint count for each species in this study and ape T2T genomes, using MFA as the reference. Consistent with T2T-CHM13-based comparisons, lemurs exhibit significantly higher breakpoint frequencies than haplorrhines when aligned to MFA (p < 0.001). The aye-aye (DMA) represents an outlier with markedly fewer breakpoints, likely due to limited alignment coverage (∼200 Mbp aligned sequence).

**Figure S4.**
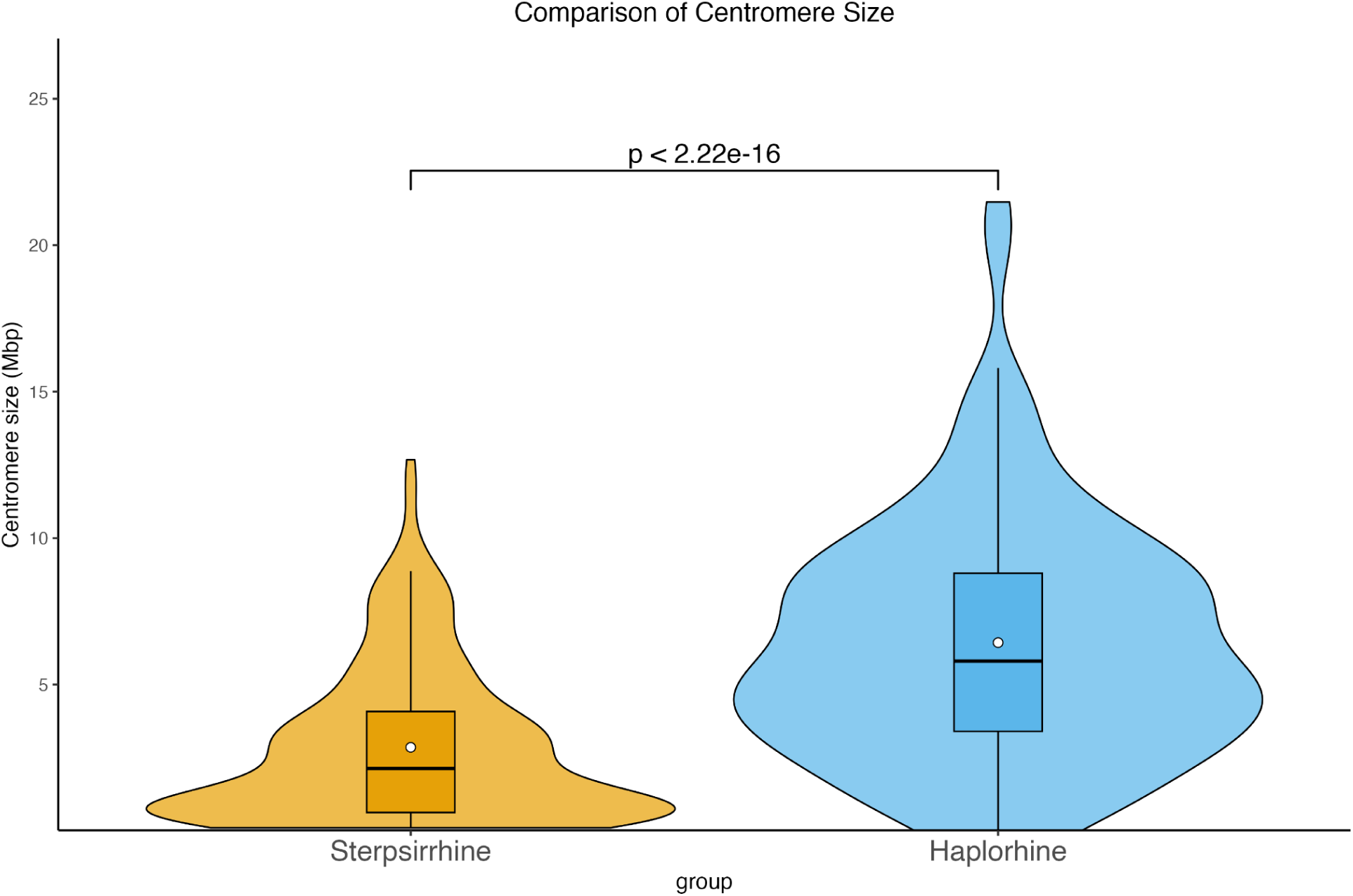
Centromere size comparison in strepsirrhines and haplorhines. We grouped the centromere sizes in two groups: strepsirrhine (eight lemur species) and haplorhine (MFA, apes and T2T-CHM13) and plotted their distributions. The difference is highly significant showing that strepsirrhine centromeres are smaller than haplorhines.

**Figure S5.**
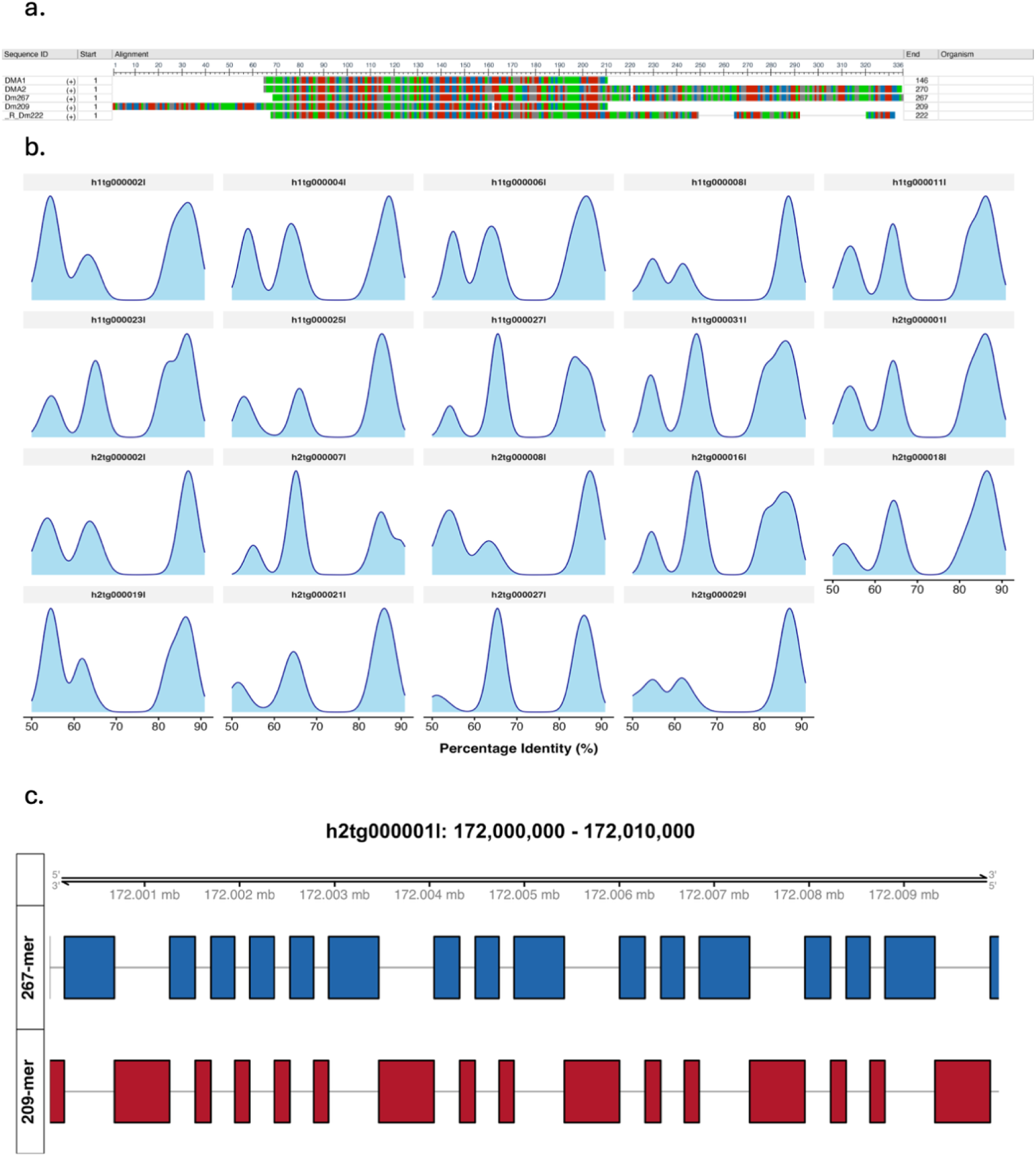
Detailed features of *Daubentonia madagascariensis* centromere. Features of DMA centromeres. a, Alignment of the three monomers found in this study along with the two monomers found in Lee et al., 2011. b, Density plots of identity of each complete centromere in the genome with respect to the most frequent canonical monomer, Dm267. c, High-resolution structure of one of the centromeres showing the repeats, Dm267 and Dm209, interspersed with each other.

**Figure S6.**
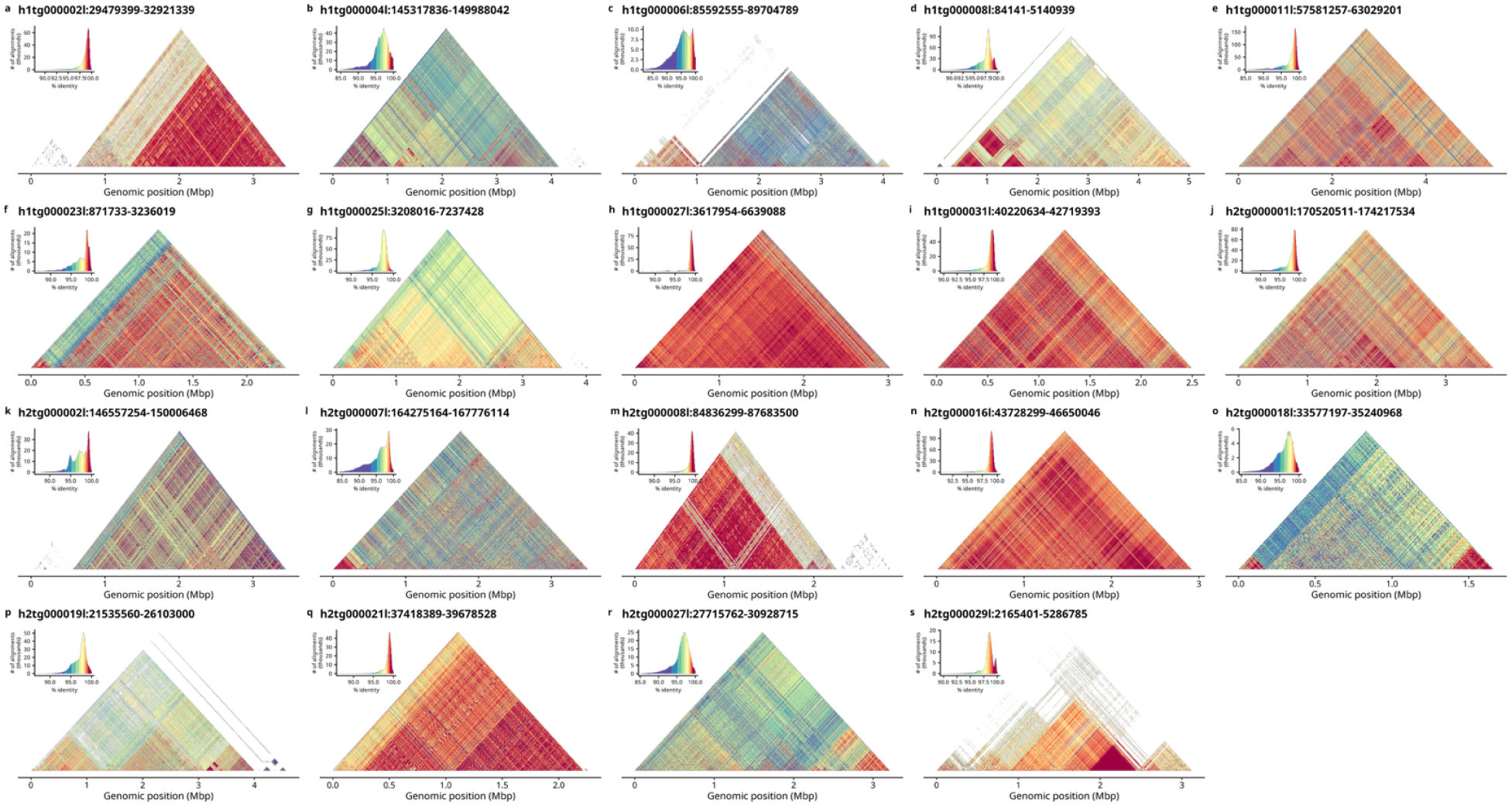
StainedGlass heatmaps for all assembled centromeres of *Daubentonia madagascariensis*.

**Figure S7.**
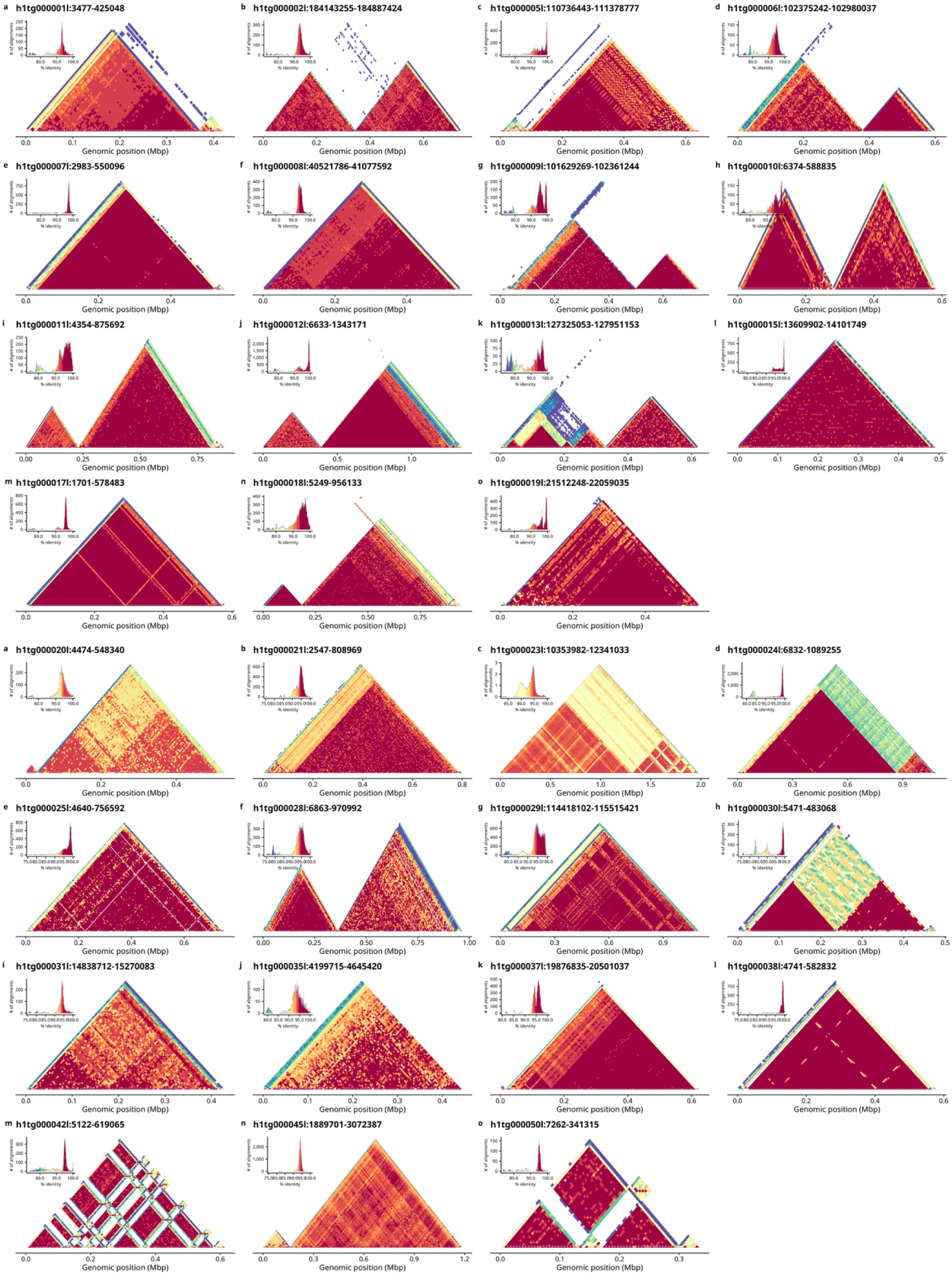
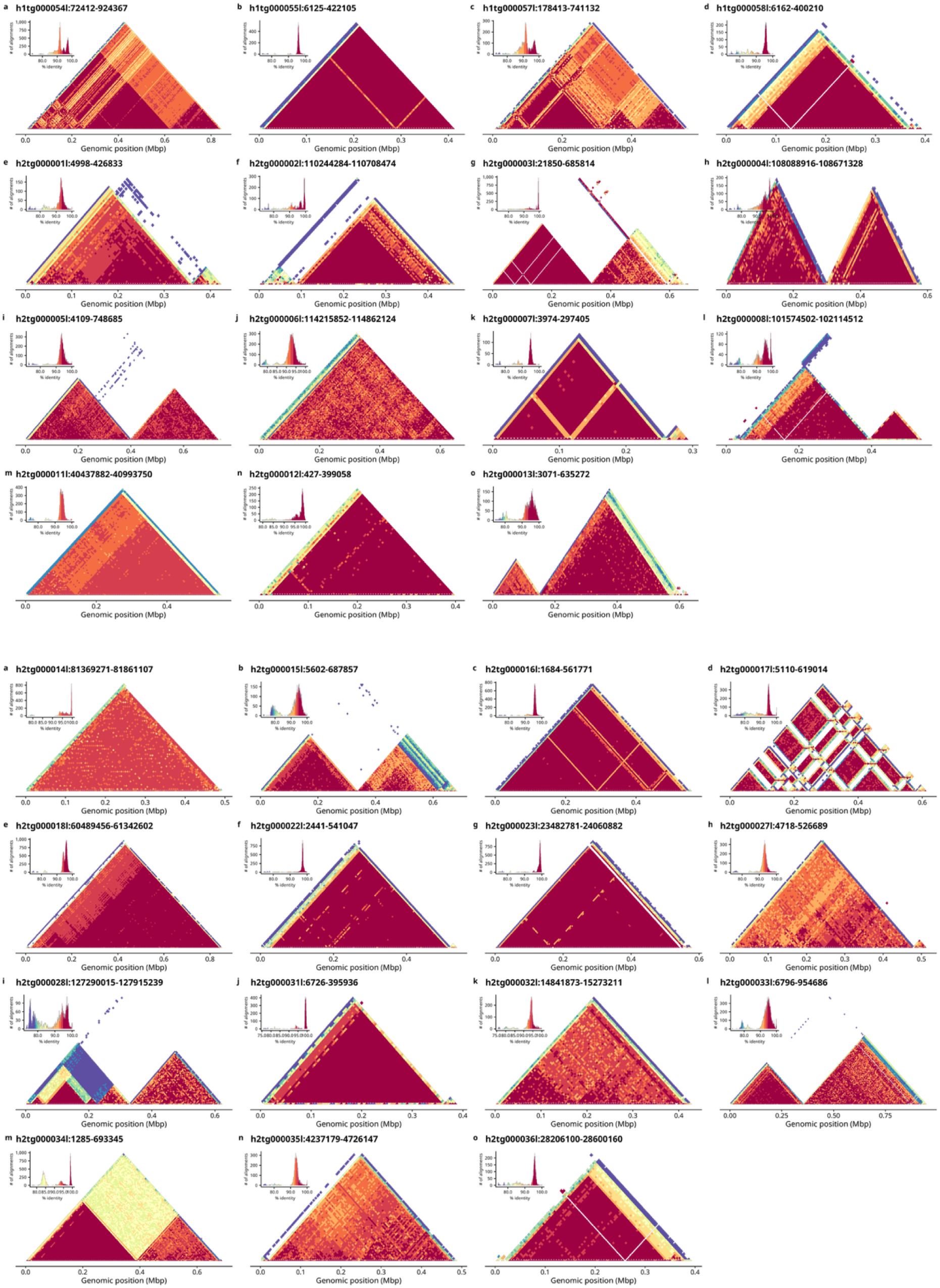
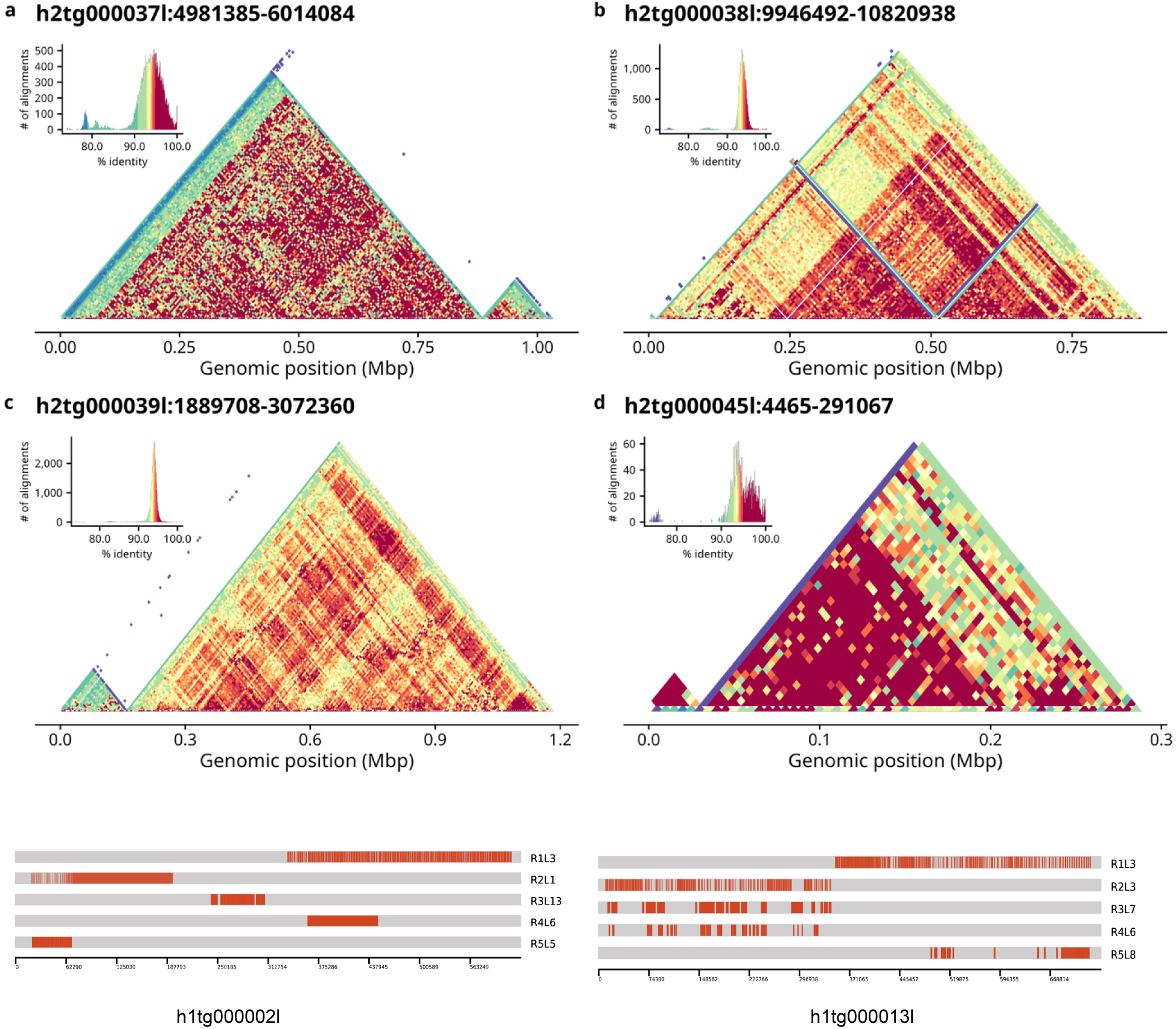
StainedGlass heatmaps for all the centromeres of *Cheirogaleus medius*. Heatmaps for centromeres of CME. The double “triangle” structure is made of two different HOR structures of the same Cm143 monomer. Examples of HOR structures for two centromeres are shown below the heatmap panel.

**Figure S8.**
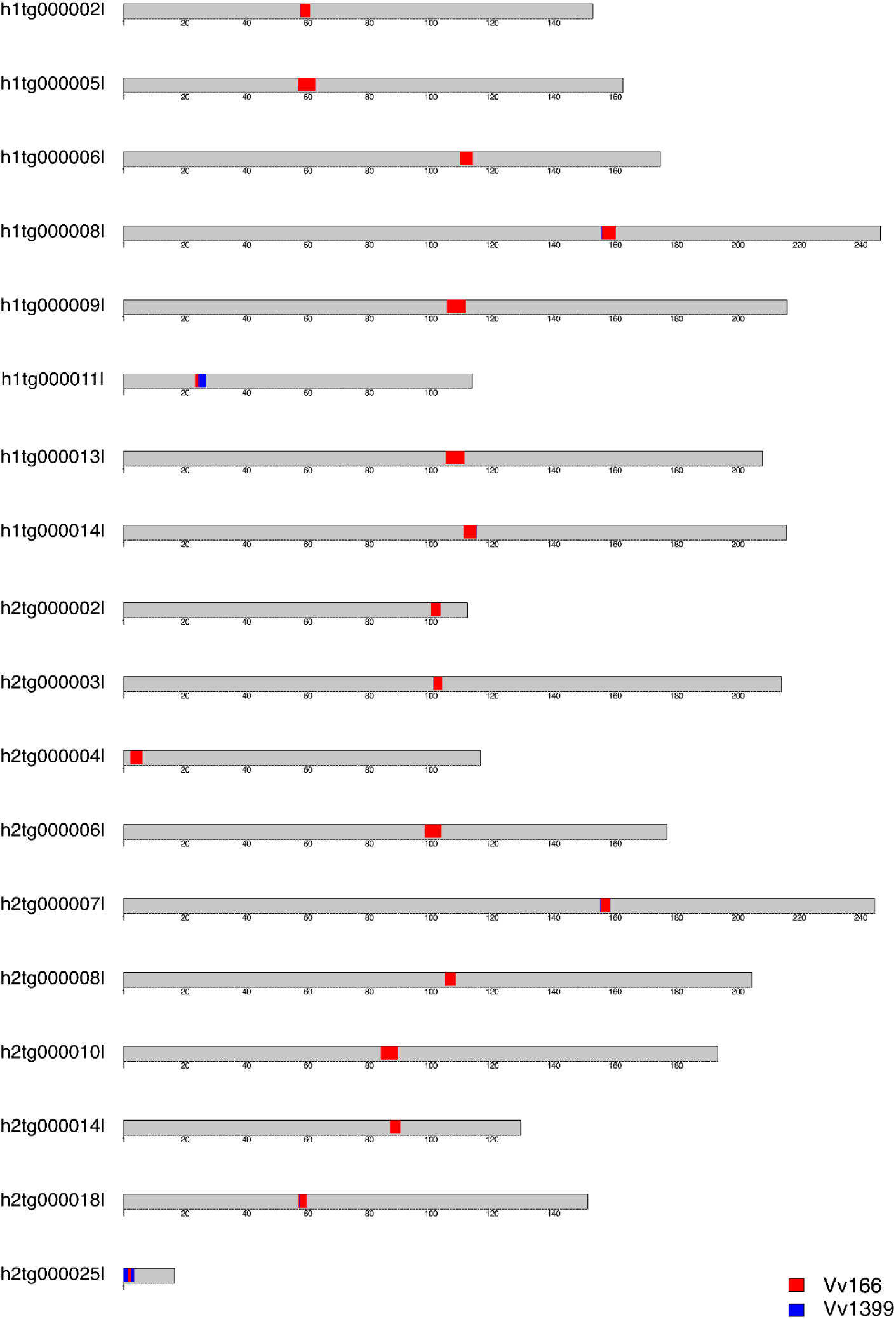
Ideogram showing centromeres on contigs of *Varecia variegata* (VVA). Ideogram showing the centromeres in each contig, with their repeat composition, whether they consist of Vv166 or Vv1405. Three types can be observed, made with either one of the two, or fused like h1tg000011l.

**Figure S9.**
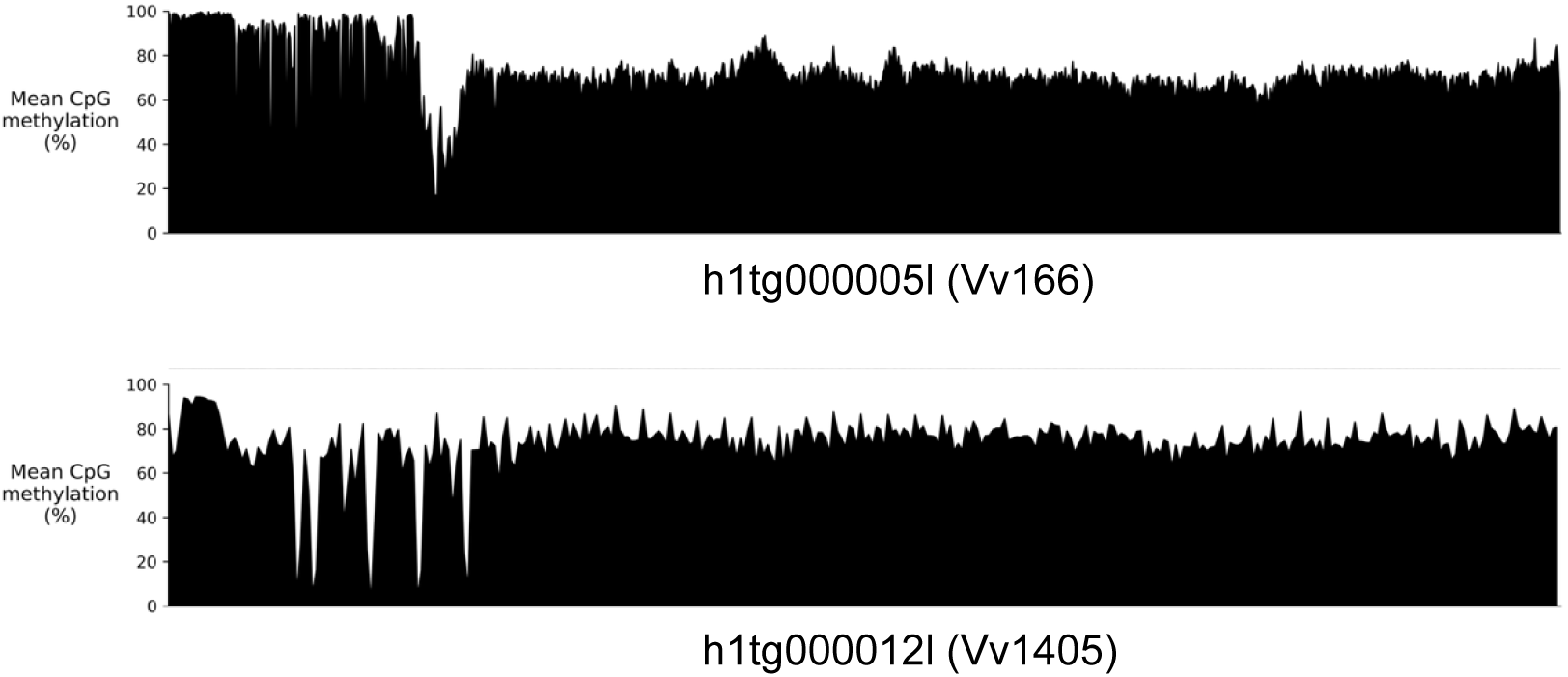
Methylation profiles of VVA centromeres consisting of different monomers. The difference in the methylation profiles of centromeres consists of Vv166 with a single CDR (above-h1tg000005l) and other with ragged CDR consisting of Vv1399 with putative small Vv166 arrays (below-h1tg000012l).

**Figure S10.**
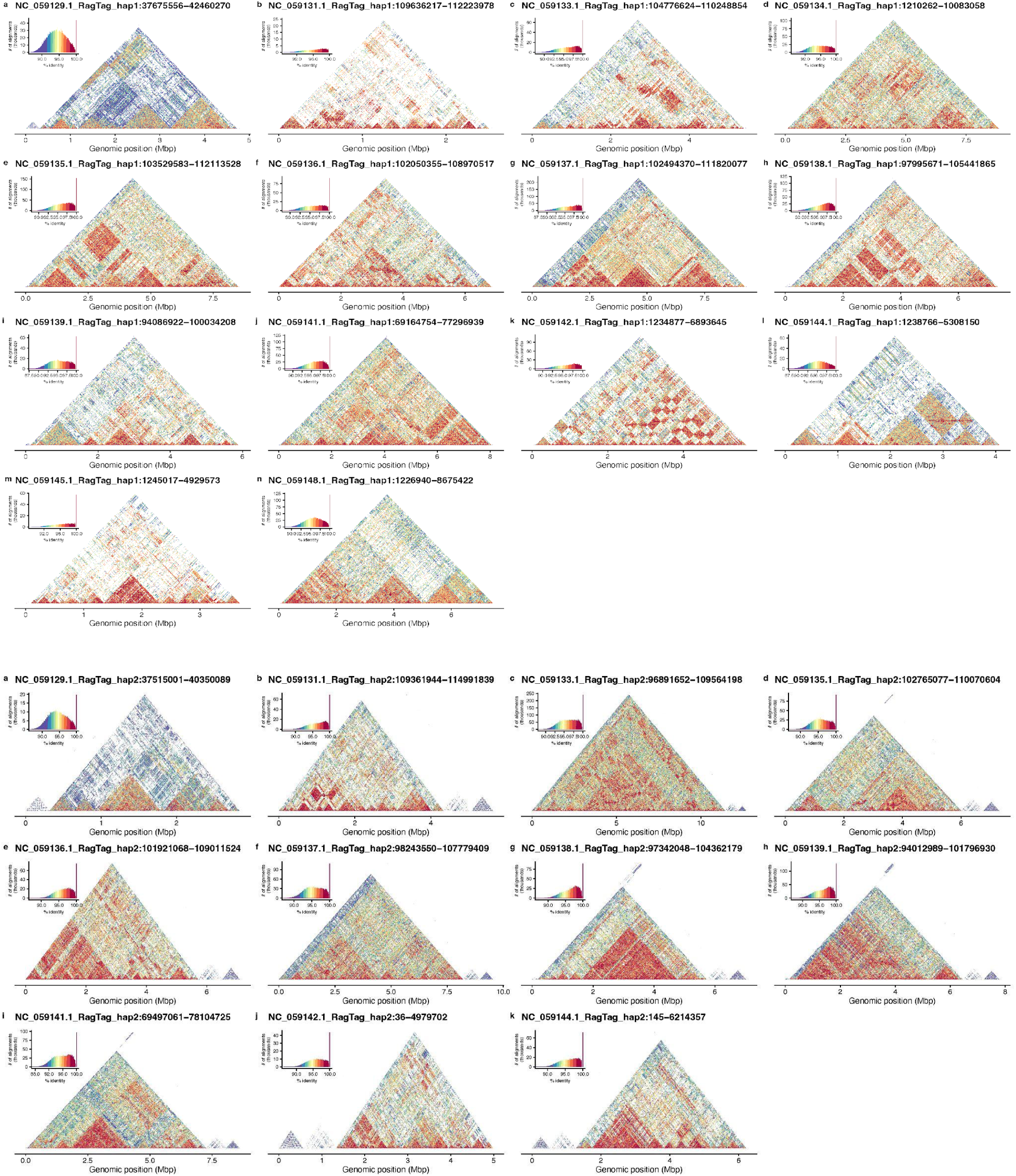
StainedGlass heatmaps for all the centromeres of *Lemur catta*. Heatmaps of all the complete centromeres of LCA. As described in the main text, the average identity of monomers of Lc41 is low due to the small size of the monomer and additionally could be because of the presence of partial polymers of 76 and 114 bp.

**Figure S11.**
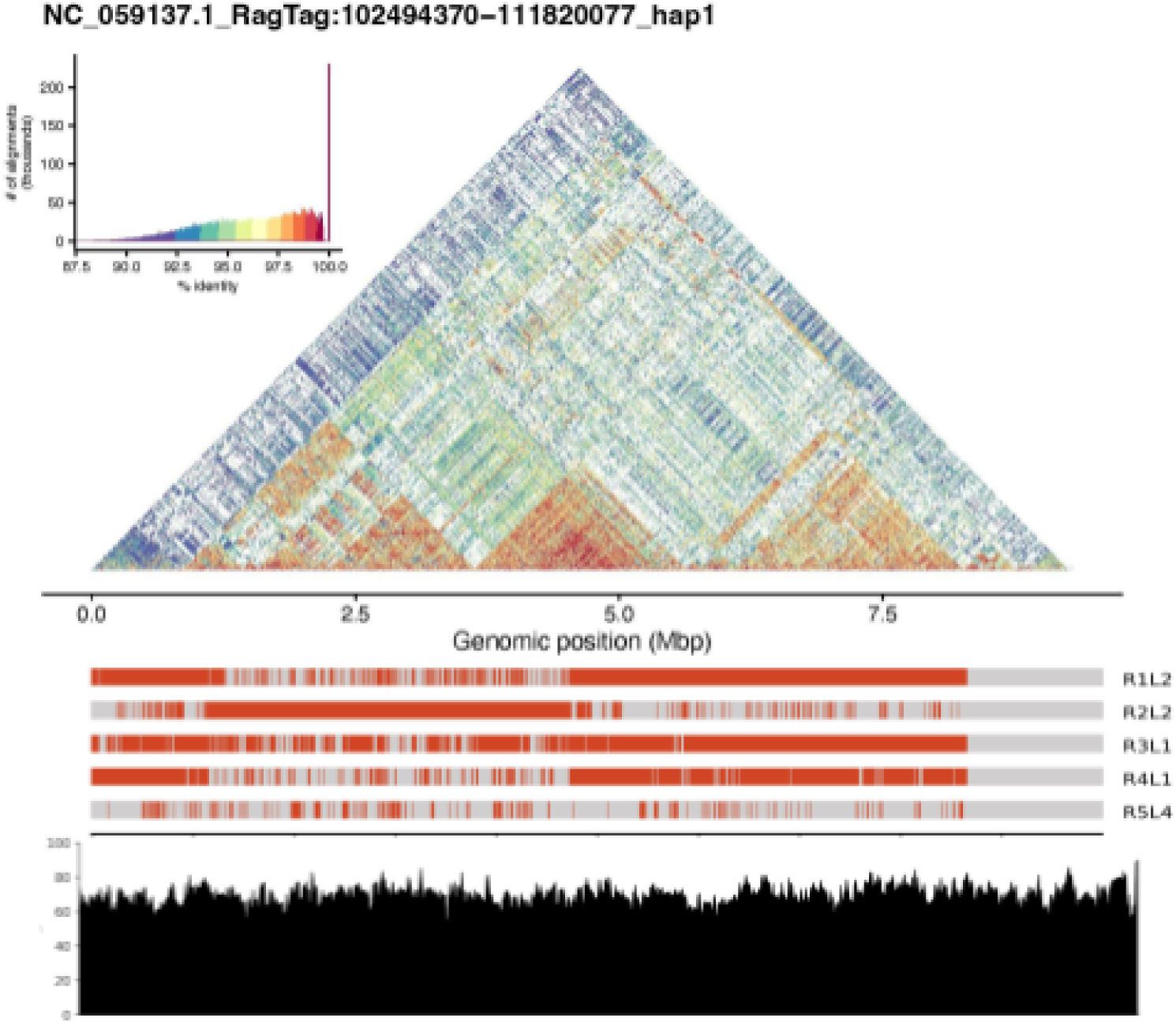
Structure of a single Lemur *catta centromere*. Here the centromere is showing an absence of a prominent CDR. Overall structure of an LCA centromere. The methylation profile does not show any prominent CDR and the whole profile is jagged.

**Figure S12.**
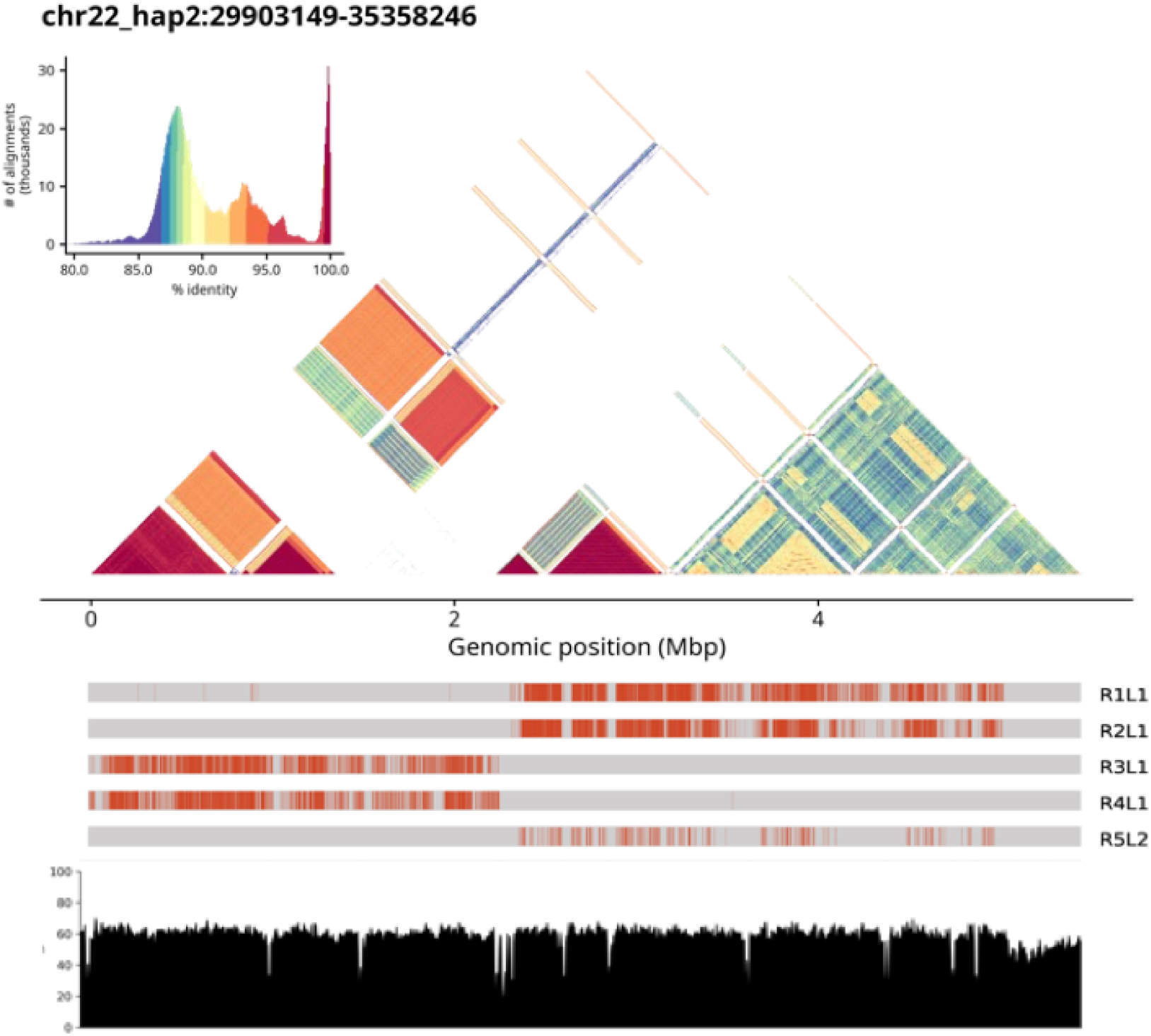
Structure of a single *Microcebus murinus* centromere. There are clearly two kinds of HORs in the centromere. They are not multi-monomeric and are only differentiated with their ranks. The methylation profile has multiple regions of hypomethylation dips.

**Figure S13.**
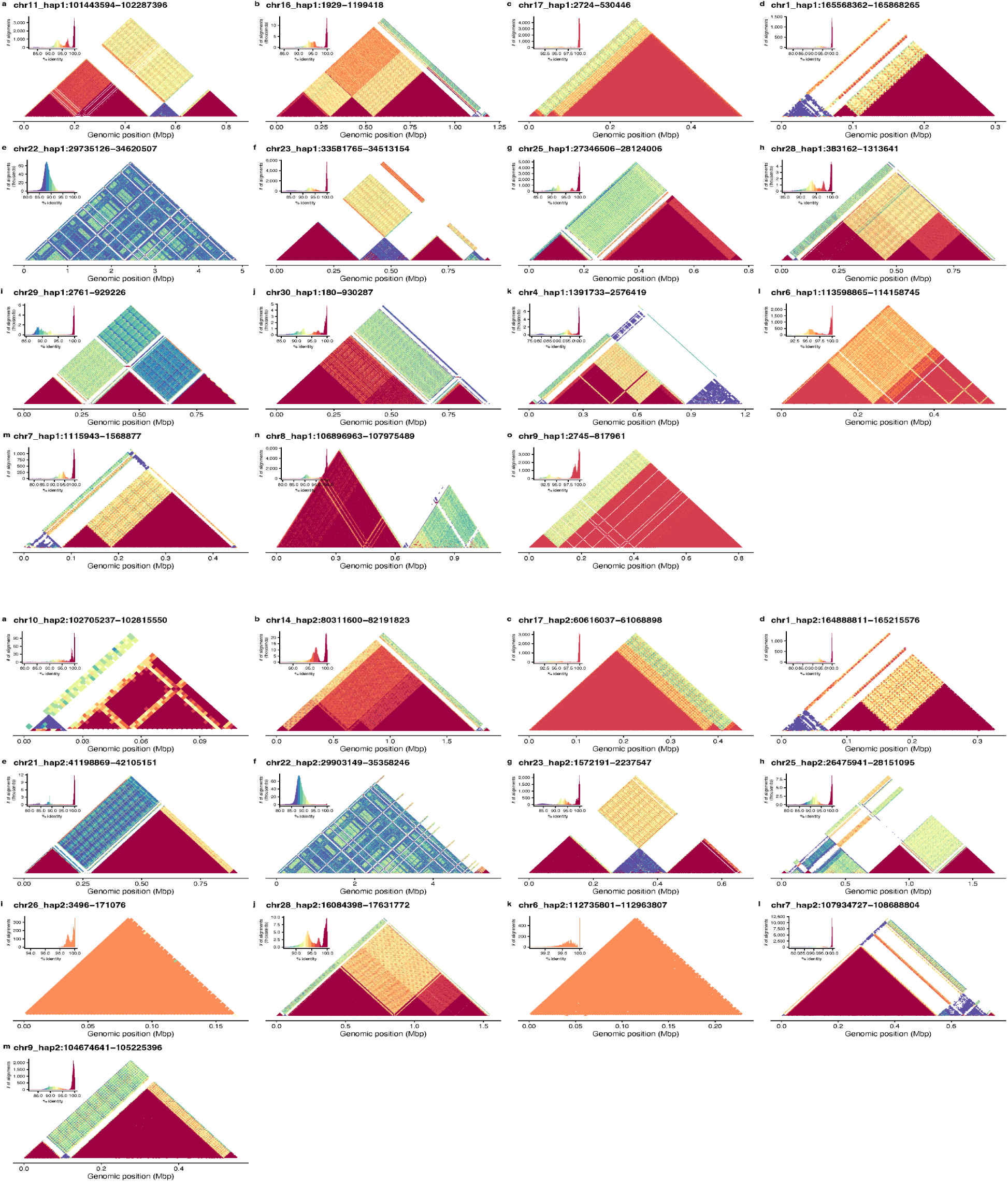
StainedGlass heatmaps for all the centromeres of *Microcebus murinus*.

**Figure S14.**
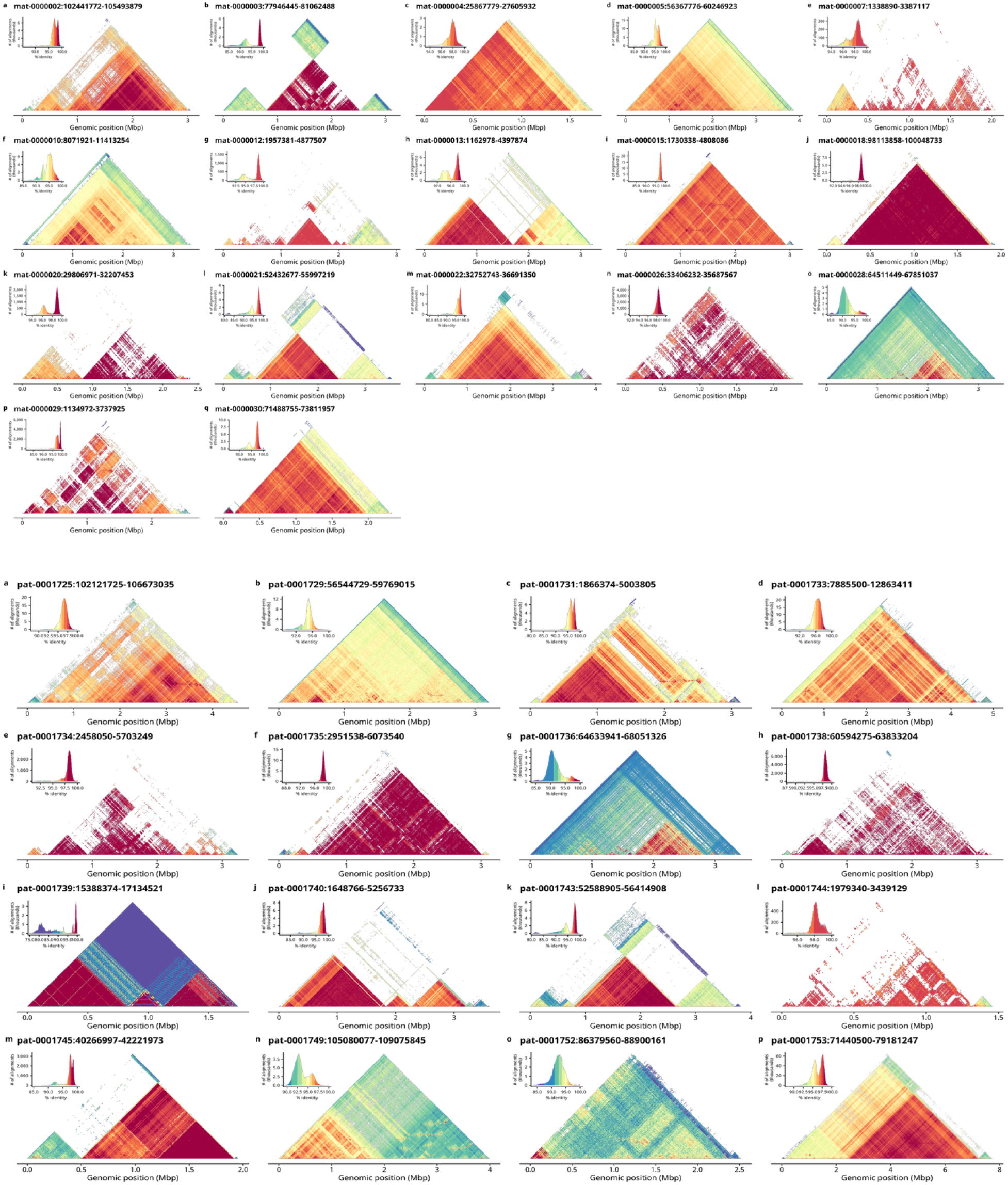
StainedGlass heatmaps for all the centromeres of *Propithecus coquereli*.

**Figure S15.**
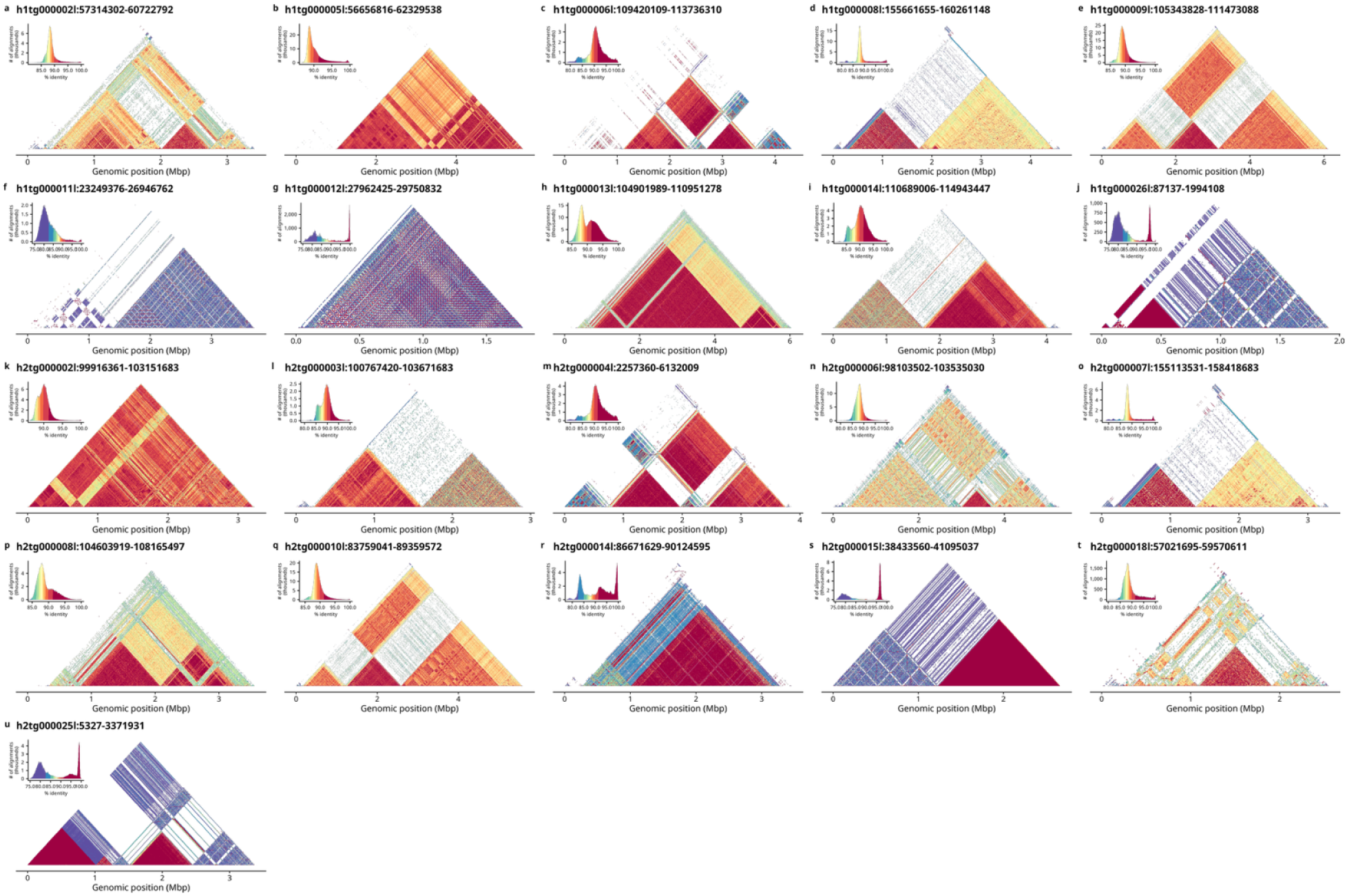
StainedGlass heatmaps for all the centromeres of *Varecia variegata*.

**Figure S16.**
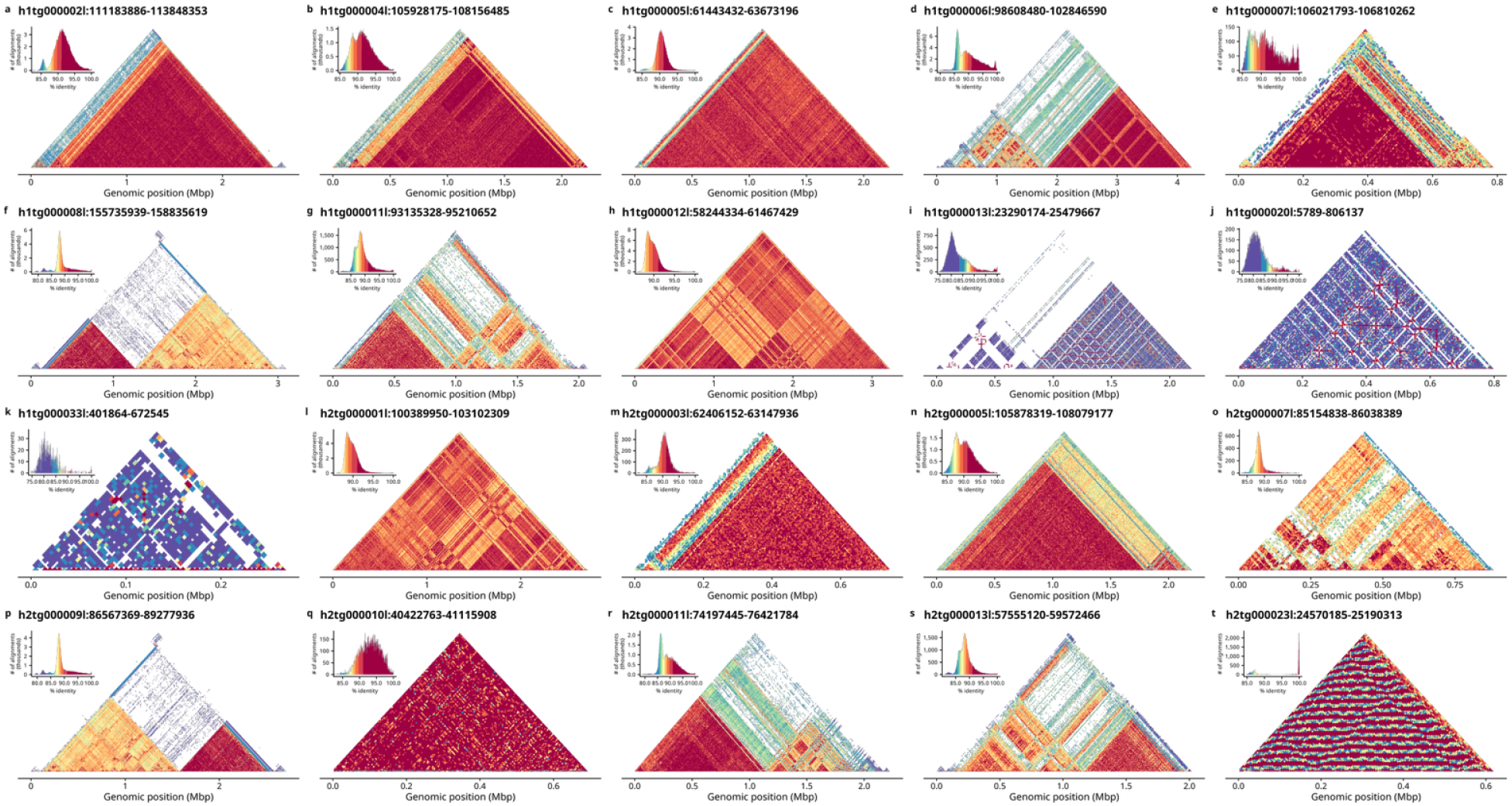
StainedGlass heatmaps for all the centromeres of *Varecia rubra*.

**Figure S17.**
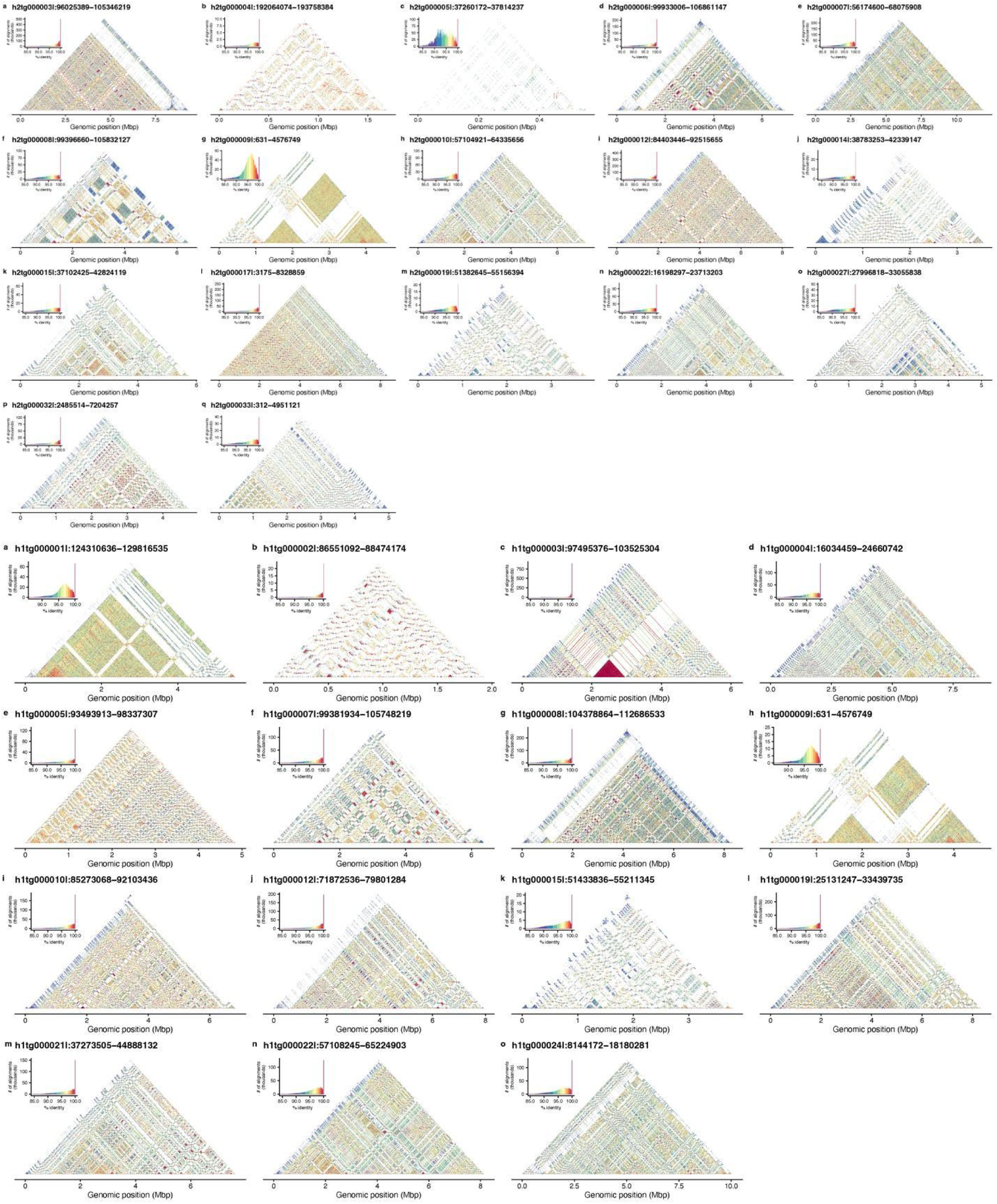
StainedGlass heatmaps for all the centromeres of *Eulemur collaris*.

**Figure S18.**
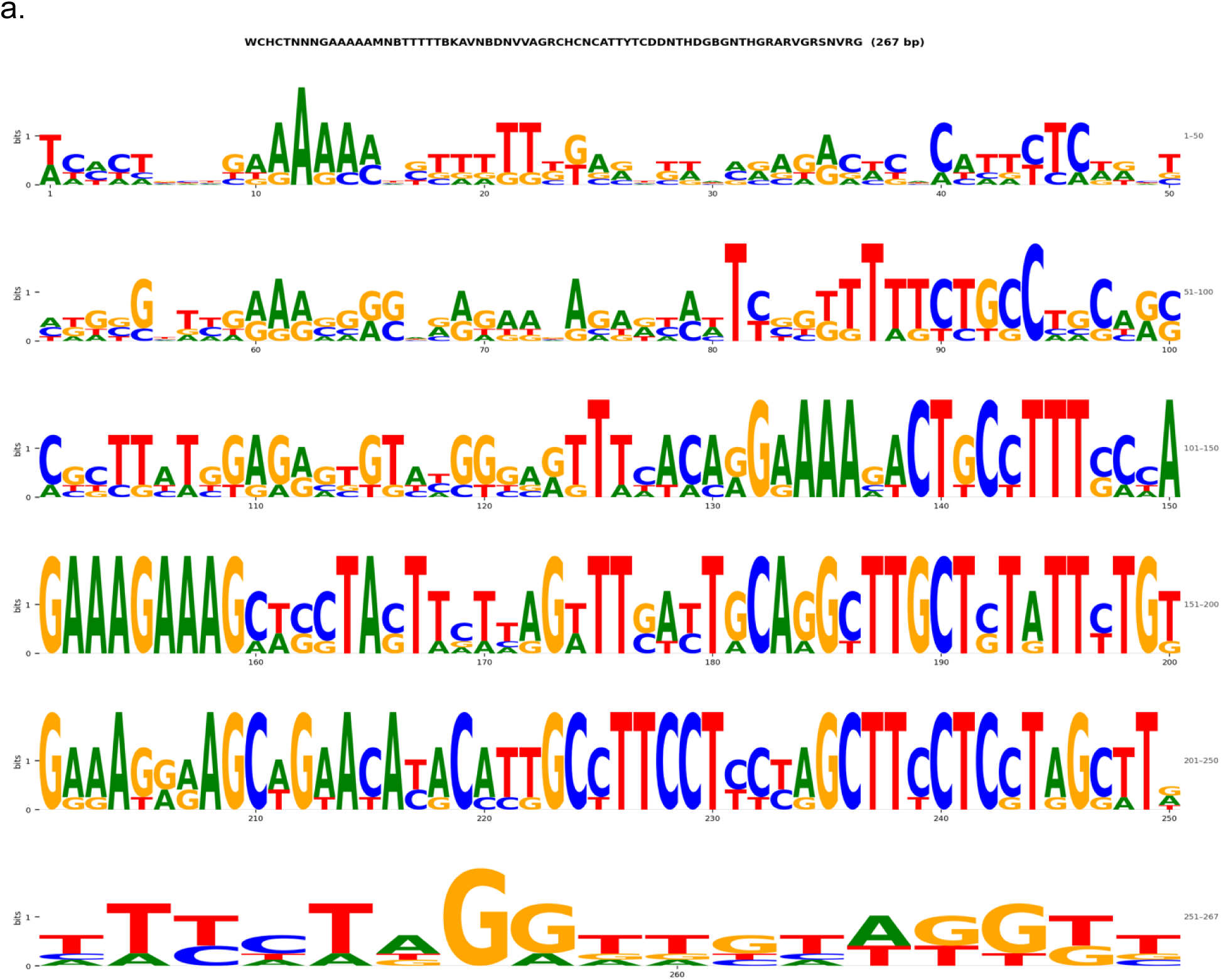
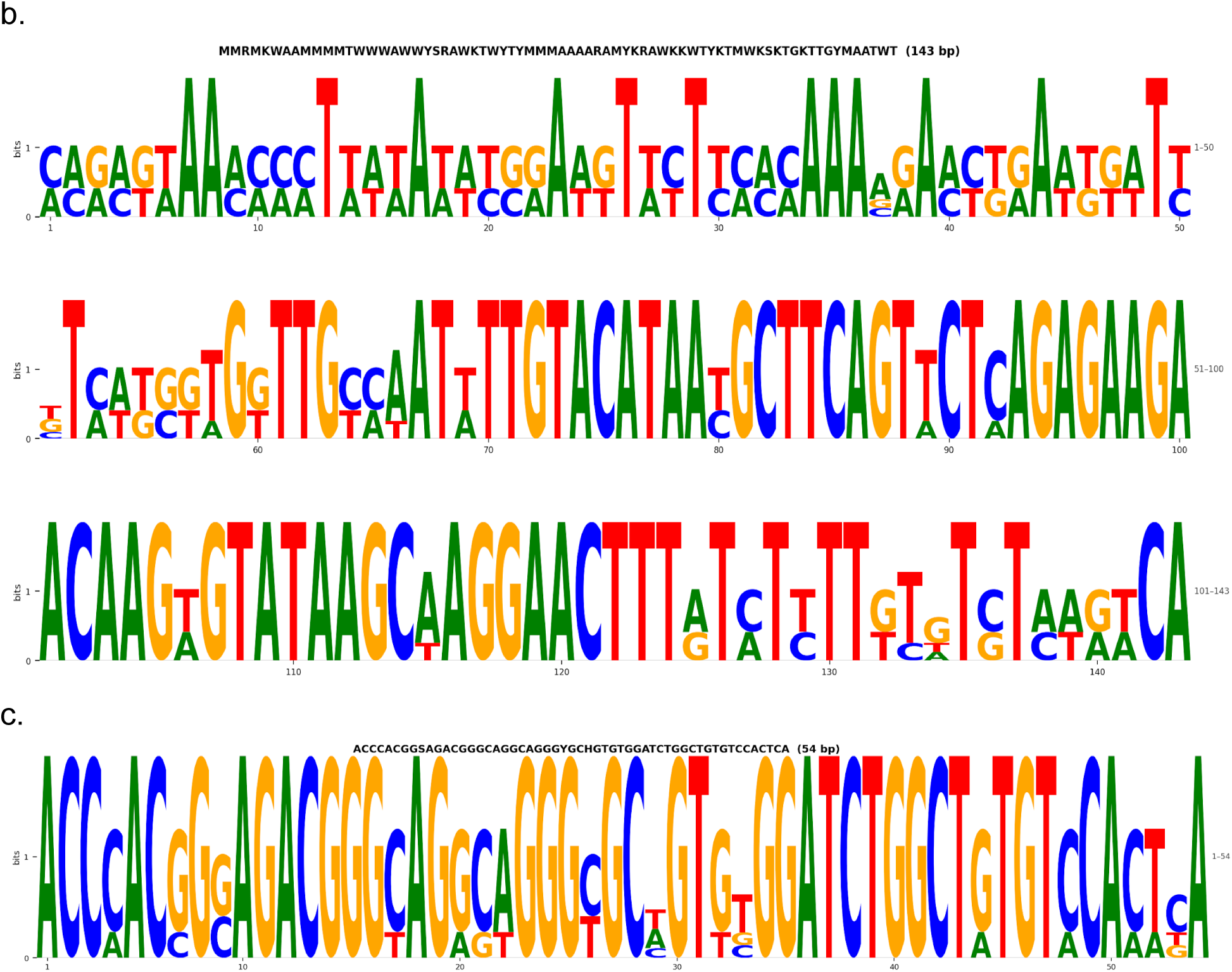
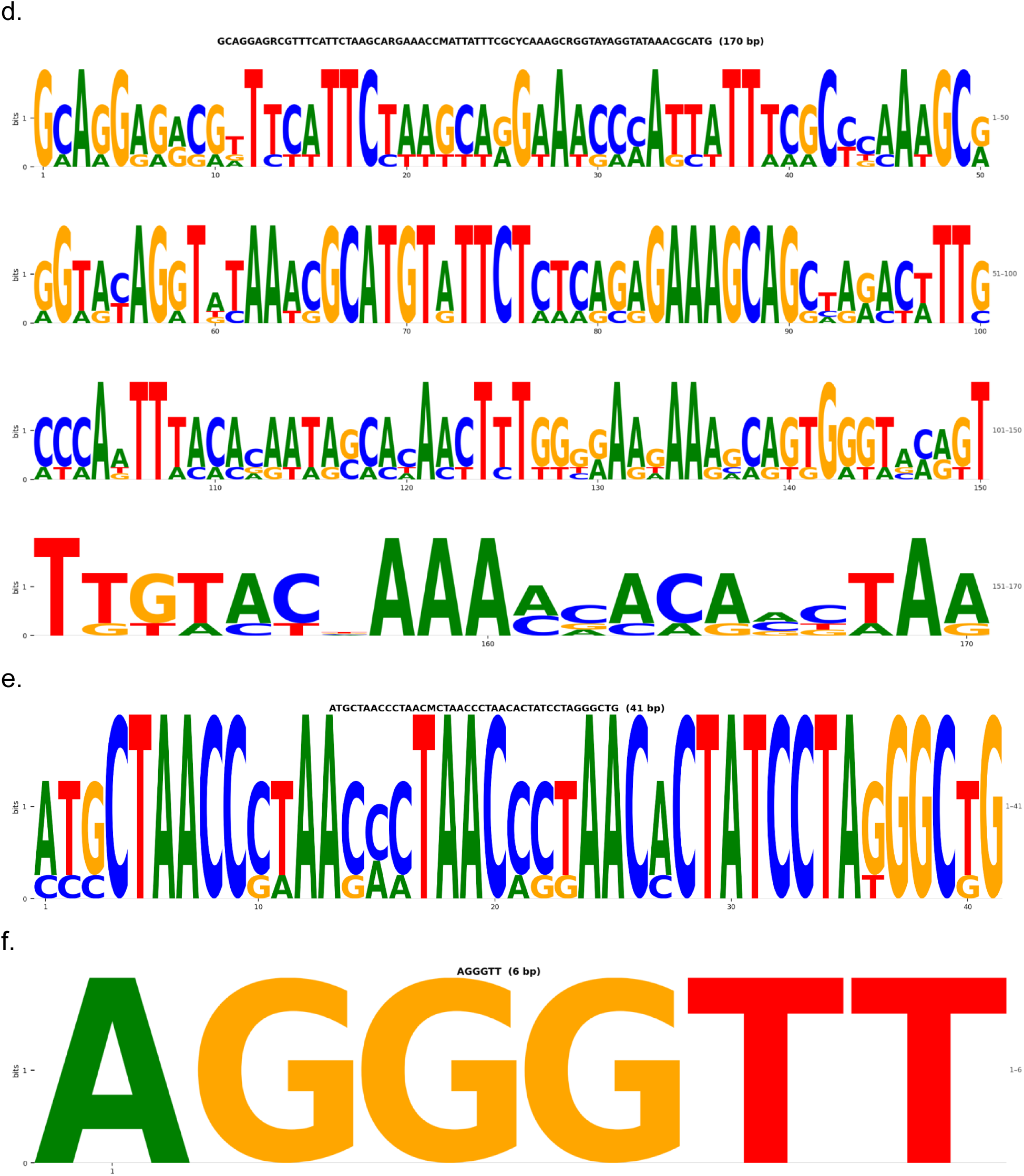
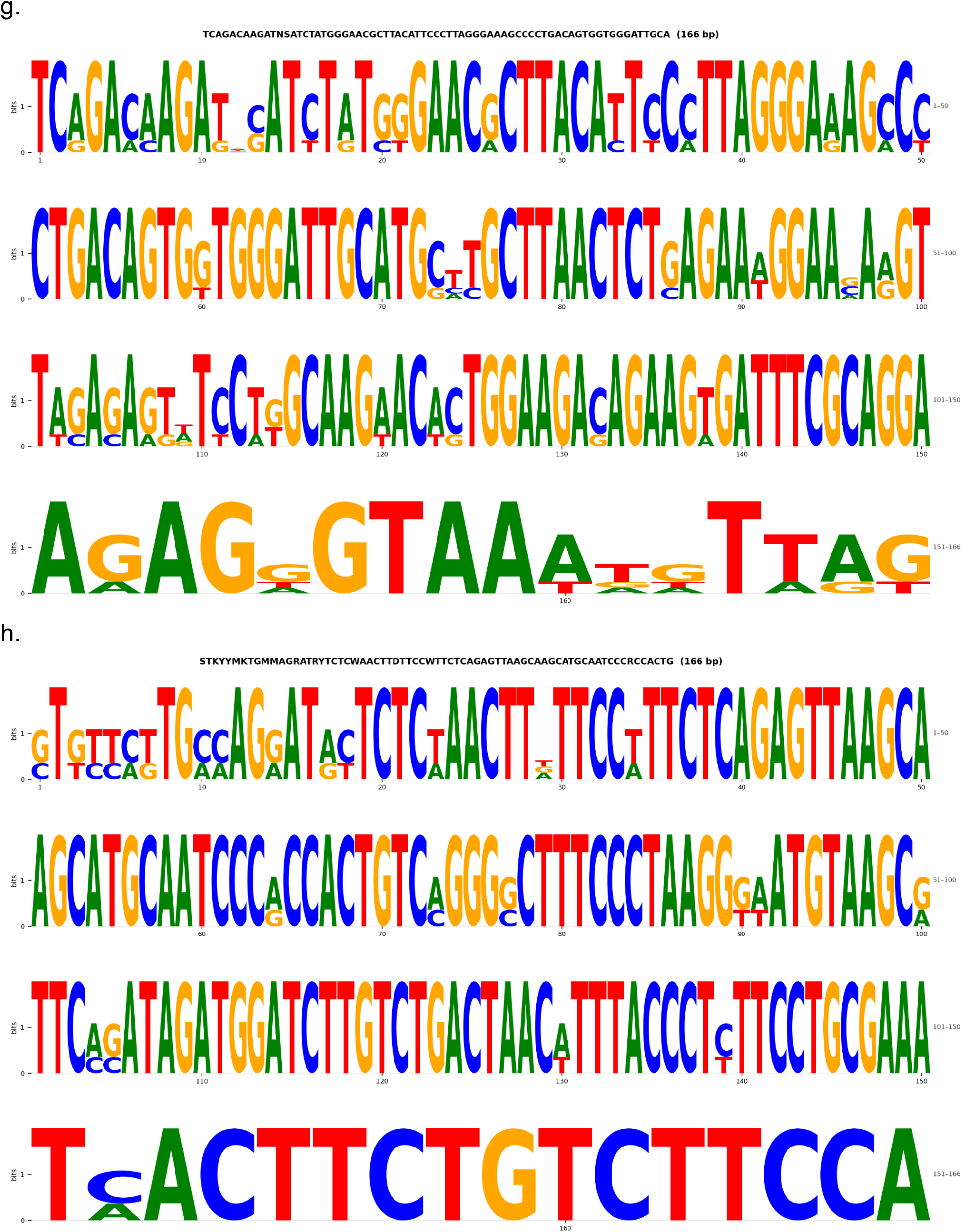
Logos plots for all the monomers across eight species of lemurs. Logos plots for the consensus sequences of the most common monomers in all the eight lemur species. The consensus was taken from the MEME suite, by giving a random 100 kbp centromere sequence and giving the exact monomer size. a, DMA; b, CME; c, MMU; d, PCO; e, LCA; f, ECO; g, VRU; h, VVA

**Figure S19.**
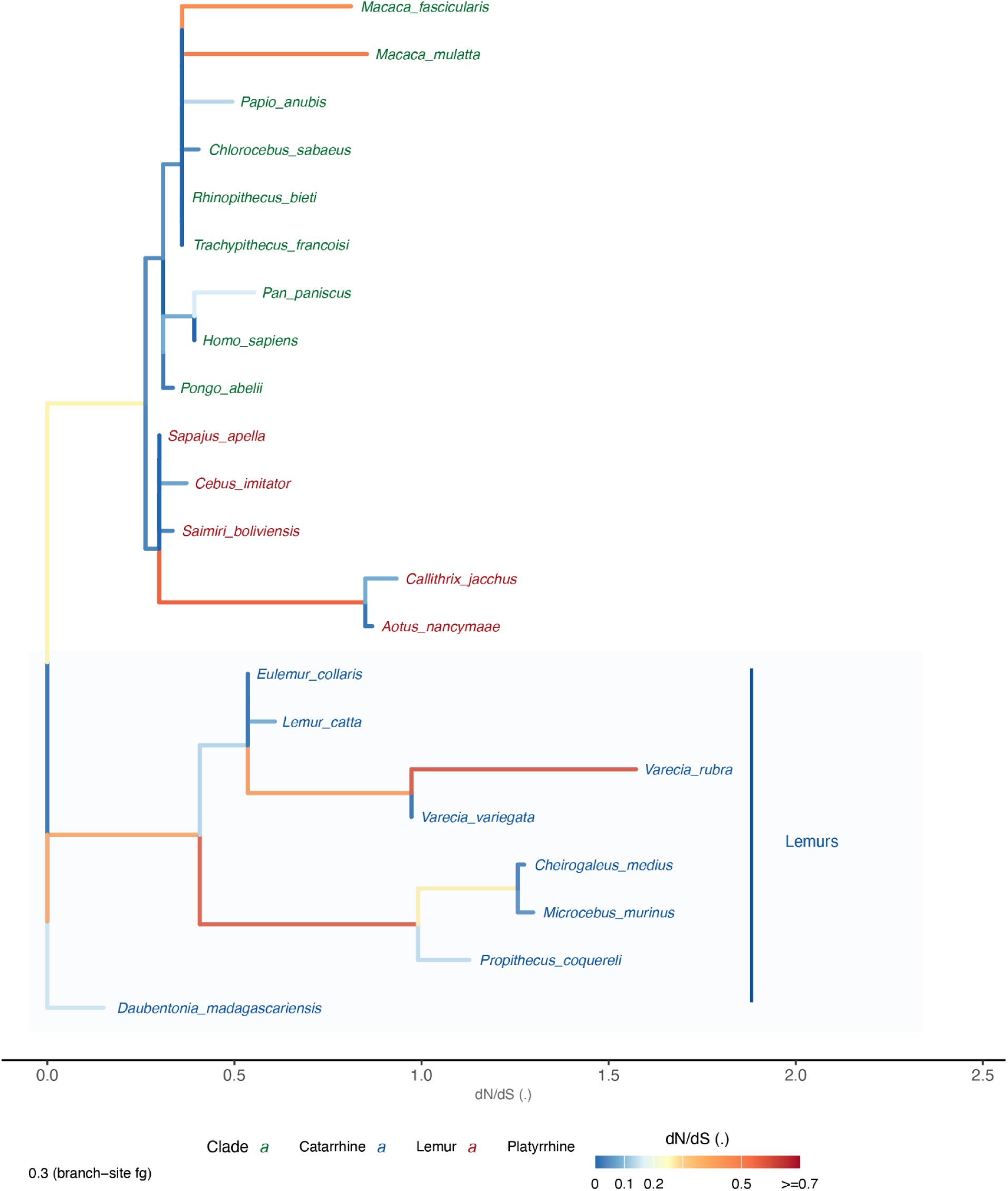
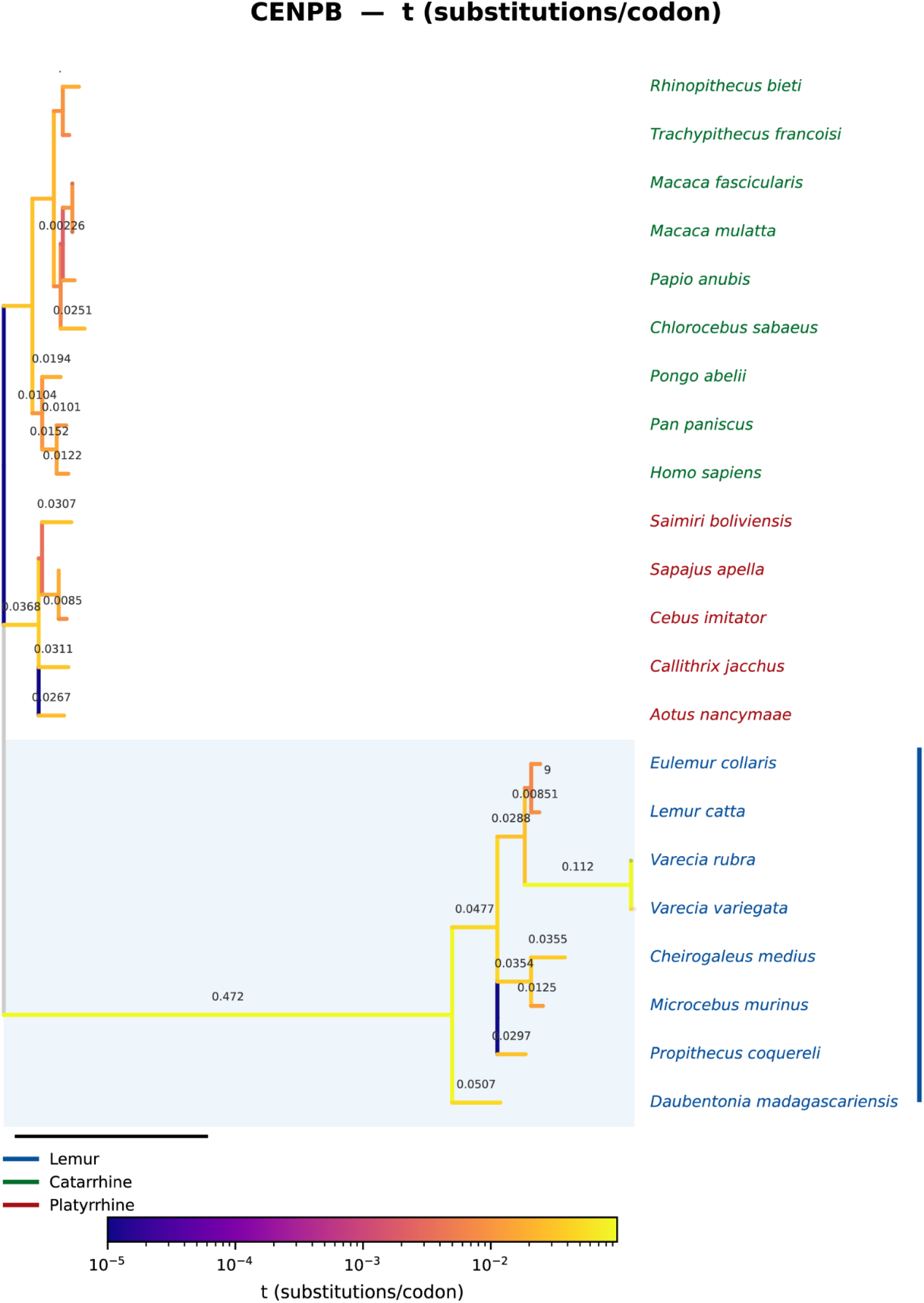
Selection on CENP-B genes is depicted with two trees. a, Branch lengths are plotted with omega values showing stronger selection in lemur clade. b. Branch length as substitutions per codon values showing the longer internal branch in lemurs.

**Figure S20.**
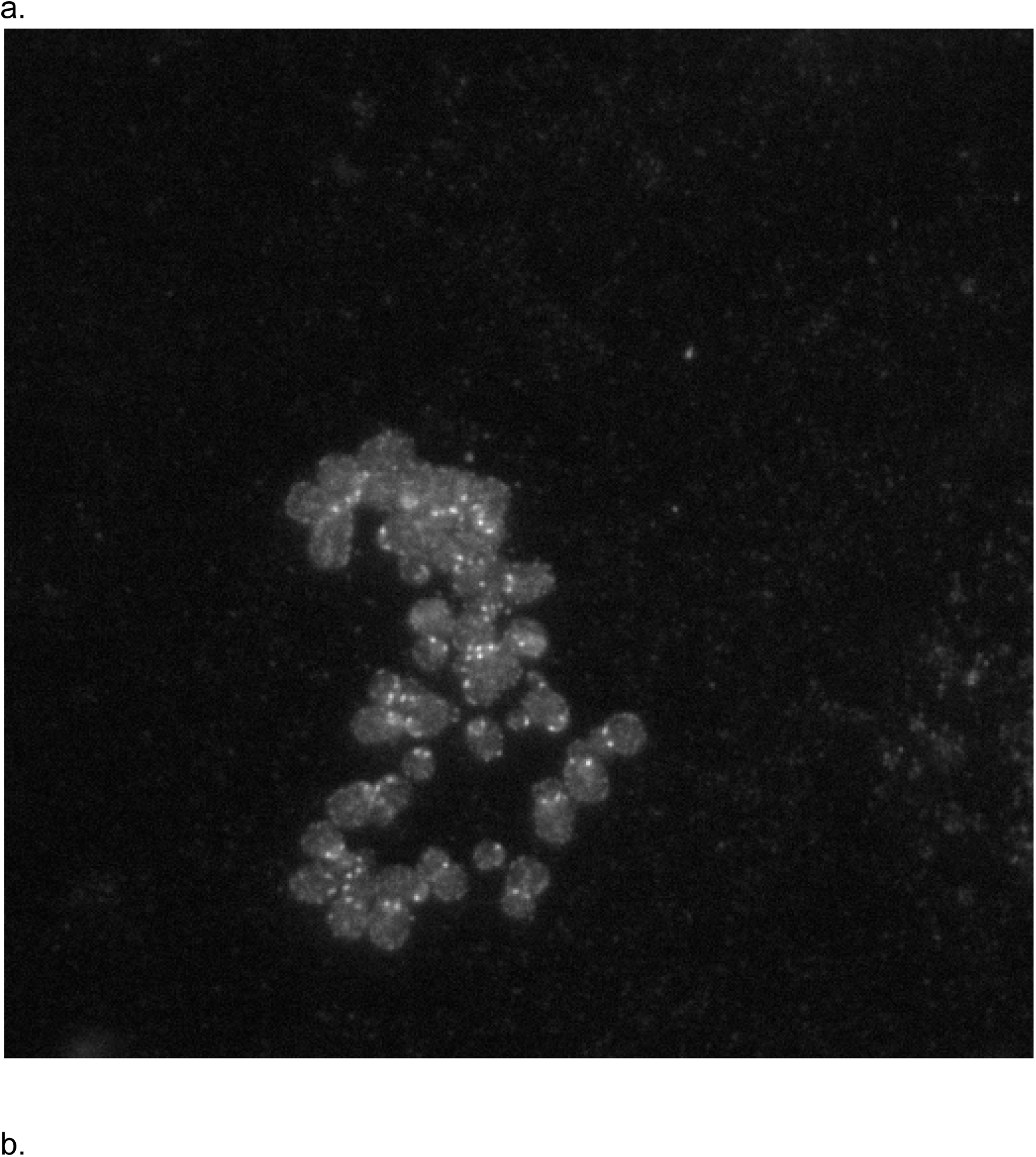
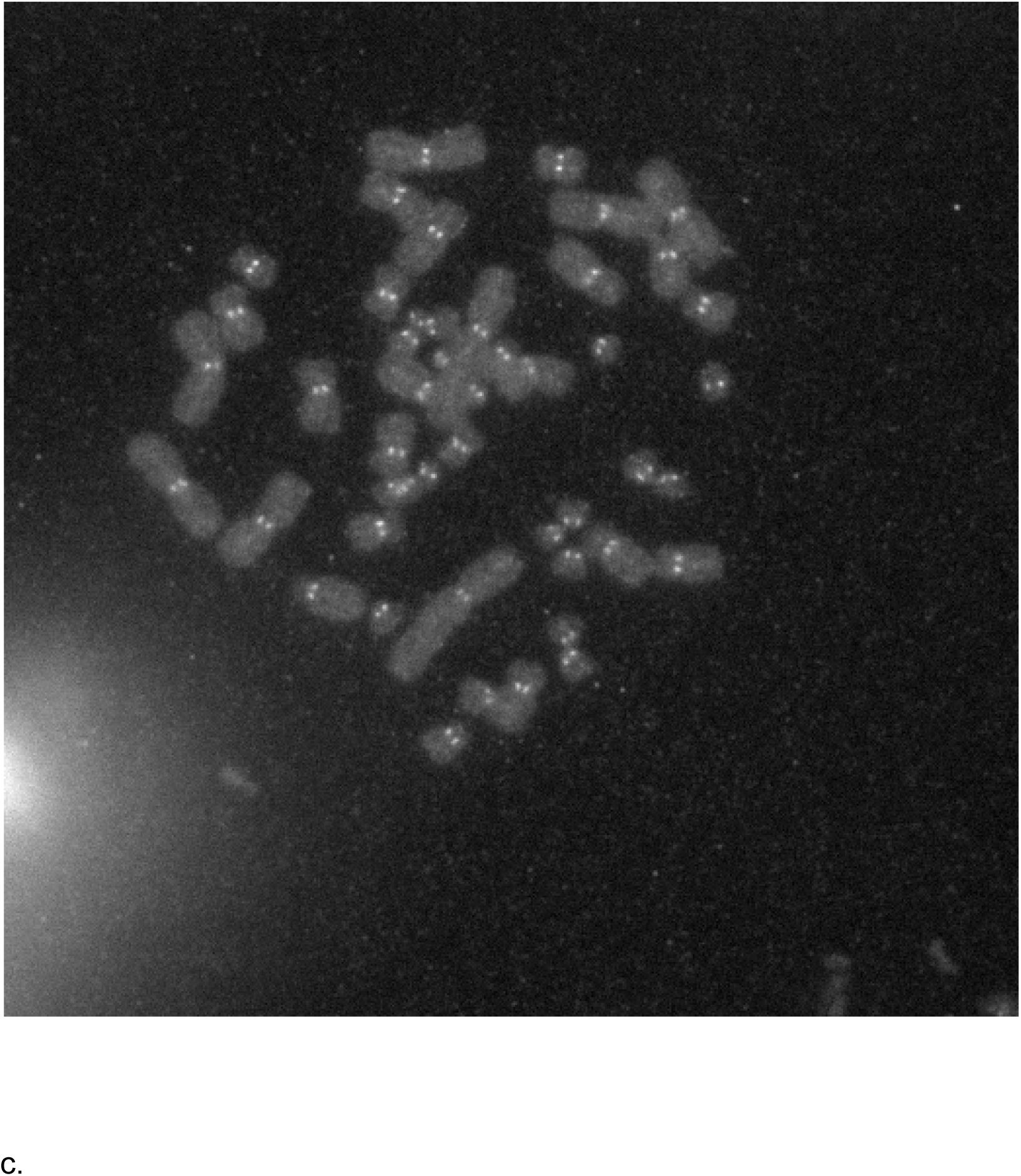
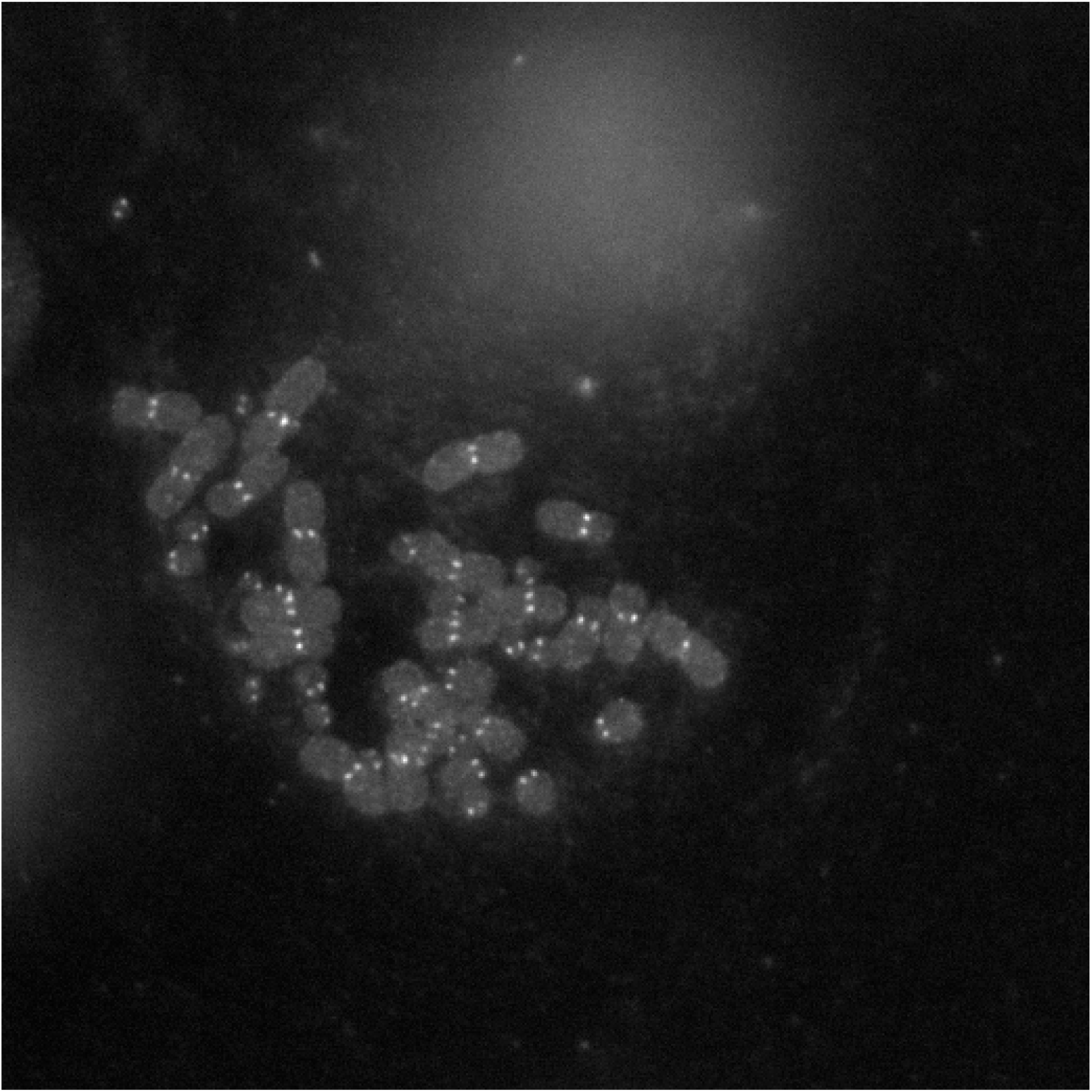
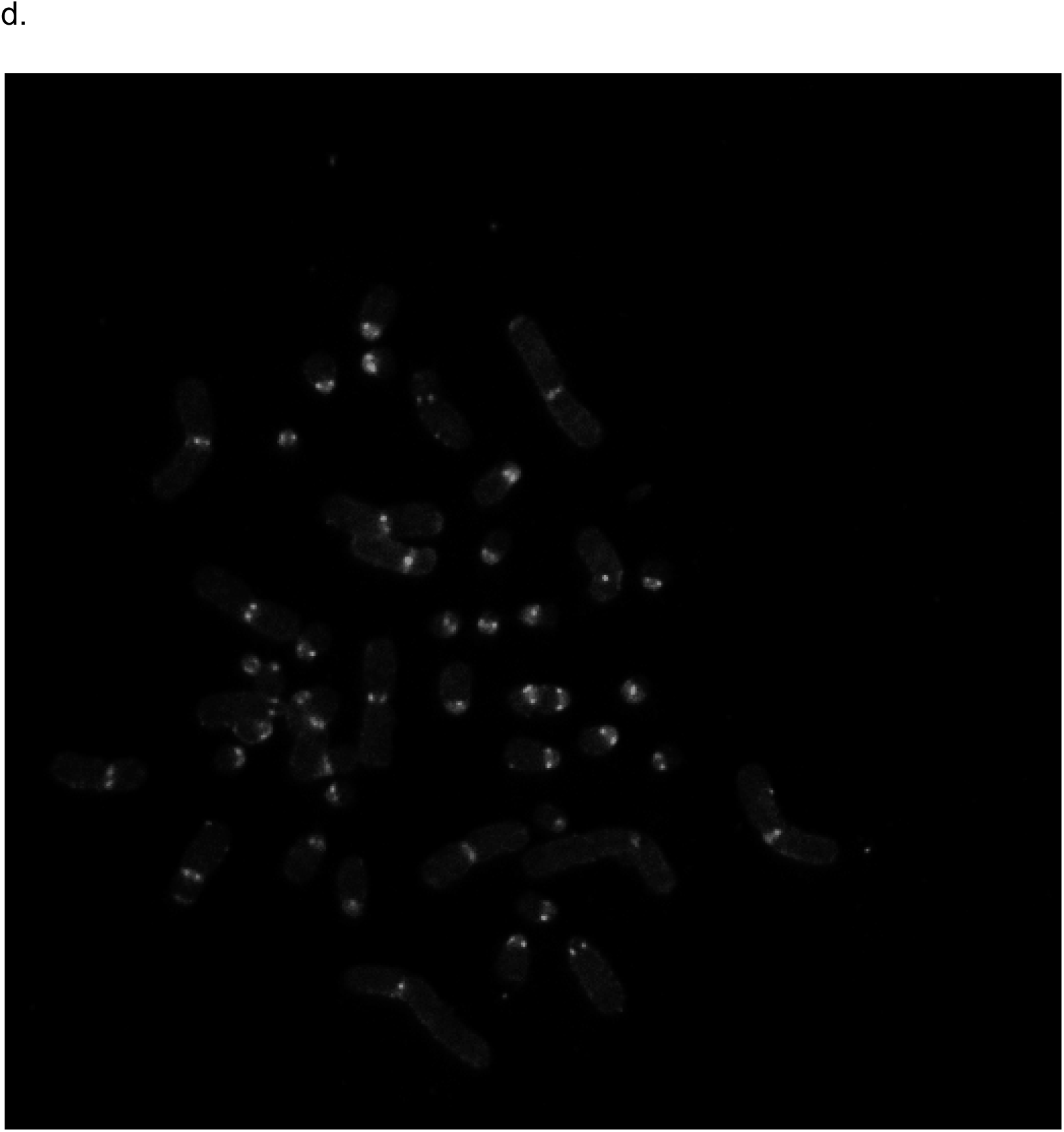
Raw FISH images of four species. Original raw images of FISH as seen in metaphase cells. a, LCA; b, PCO; c, VVA; d, ECI.

